# Intracellular amorphous calcium carbonate biomineralization in methanotrophic gammaproteobacteria was acquired by horizontal gene transfer from cyanobacteria

**DOI:** 10.64898/2025.12.07.689555

**Authors:** Benzerara Karim, Millet Maxime, Skouri-Panet Fériel, Gaschignard Geoffroy, Mehta Neha, Bezard Malo, Caumes Geraldine, Daniel M. Chevrier, Dezi Manuela, Duverger Arnaud, Guigner Jean-Michel, Gutiérrez-Preciado Ana, Christopher T. Lefevre, López-García Purificación, Menguy Nicolas, Caroline L. Monteil, Pehau-Arnaudet Gerard, Penard Esthel, Pereiro Eva, Scandola Cyril, Travert Cynthia, Vantelon Delphine, Duprat Elodie, Callebaut Isabelle, Moreira David

**Affiliations:** Sorbonne Université, Muséum National d’Histoire Naturelle, UMR CNRS 7590. Institut de Minéralogie, de Physique des Matériaux et de Cosmochimie (IMPMC), 75005 Paris, France; Aix-Marseille Université, CEA, CNRS, BIAM, UMR7265, Institut de Biosciences et Biotechnologies Aix-Marseille, CEA Cadarache, F-13115, Saint-Paul-lez-Durance, France; Unité d’Ecologie Société et Evolution, CNRS, Université Paris-Saclay, AgroParisTech, Gif-sur-Yvette, France; Institut Pasteur, Université Paris Cité, Ultrastructural Bioimaging Unit, 75015 Paris, France; ALBA Synchrotron Light Source, Cerdanyola del Vallés, Barcelona 08290, Spain; Synchrotron SOLEIL, L’Orme des Merisiers, 91190 Saint-Aubin, France

**Keywords:** Amorphous calcium carbonate, methanotrophs, Cyanobacteria, biomineralization, horizontal gene transfer

## Abstract

Some bacteria genetically control the biomineralization of intracellular amorphous calcium carbonates (iACC) with potential implications for microbial physiology, evolution, bioremediation and biogeochemical cycling. Until now, this capacity has been documented in *Cyanobacteria*, the giant gammaproteobacterium *Achromatium* and a few magnetotactic *Pseudomonadota* and *Nitrospirota*. Here, we report the discovery of iACC biomineralization in members of the *Methylococcaceae*, a family of aerobic methanotrophic *Gammaproteobacteria*. A homolog of the *ccyA* gene, previously considered a diagnostic marker for iACC formation in *Cyanobacteria*, was identified in several *Methylococcaceae* genomes, based on a search of the conserved C-terminal (GlyZip)3 domain of the encoded calcyanin protein, with a sequence coverage higher than 60% and an E-value lower than 1e^-20^. Moreover, two cultivated strains, *Methylococcus geothermalis* and *Methylococcus mesophilus*, whose genomes contained the *ccyA* gene, were consistently shown to form iACC granules. The *ccyA* genes of *Methylococcaceae* and *Microcystis* share higher sequence similarity (47%) than with other *Cyanobacteria* (around 30%) within their common (GlyZip)_3_ domain, suggesting horizontal gene transfer (HGT) from an ancestral *Microcystis*-like cyanobacterium to *Methylococcaceae*. This finding extends the known taxonomic distribution of *ccyA* and suggests that the capability to biomineralize iACC was acquired by HGT, possibly in environments such as those close to the oxyclines of lakes, where *Cyanobacteria* and *Methylococcaceae* commonly co-exist. The discovery of iACC in methane-oxidizing *Methylococcaceae* highlights a previously unrecognized coupling between calcium carbonate biomineralization and methane cycling in aquatic environments, suggesting that iACC formation may play an overlooked role in microbial carbon storage and local geochemical regulation.

## 1. Introduction

Many bacteria are capable of forming a wide range of mineral phases through a process known as biomineralization. This process may serve specific biological functions such as protection, detoxification, assistance in cell movements or storage (Cosmidis and Benzerara, 2022) and has a significant impact on both local and global geochemical cycles, inspiring a variety of applications (Qin et al., 2020; Wan et al., 2024; Zhang et al., 2023). Among biomineralizing bacteria, several phylogenetically diverse species have been shown to form intracellular amorphous calcium carbonates (iACC). The giant gammaproteobacterium *Achromatium oxaliferum* was the first such organism discovered, as early as the late 19^th^ century (Schewiakoff, 1893). For a long time, it remained the only known iACC-biomineralizing bacterium (Benzerara et al., 2021; Gray, 2006). However, since 2012, diverse cyanobacteria have also been found to form iACC (Bacchetta et al., 2024; Benzerara et al., 2014, 2022; Couradeau et al., 2012). More recently, additional members of the *Pseudomonadota* and *Nitrospirota*, including magnetotactic bacteria capable of biomineralizing intracellular magnetite crystals (Fe(II)-Fe(III) oxides), have been shown to form iACC as well (Liu et al., 2021, 2025; Mangin et al., 2025; Monteil et al., 2020). This rapidly expanding diversity of iACC-forming bacteria suggests that many more such organisms likely remain to be discovered.

Forming iACC granules incurs an energy cost to cells, at least under certain conditions (Cam et al., 2017), suggesting that these granules likely confer adaptive benefits to the cells. However, their exact biological role remains poorly understood (Cosmidis and Benzerara, 2022). Some studies have proposed that iACC granules may buffer intracellular pH fluctuations, serve as carbon reservoirs, or act as ballast for the cells (Görgen et al., 2020; Monteil et al., 2020). Regardless of their specific function, iACC granules involve a particular Ca homeostasis in the cells that form them, including Ca exchange between the cells and the surrounding solution with a net inward influx (Benzerara et al., 2023; Mehta et al., 2023, 2024). Studying these granules offers valuable insights: (i) Amorphous calcium carbonate (ACC) is essential to many CaCO₃-biomineralizing eukaryotes (Gilbert et al., 2022), and the ability to form iACC may have originated early in evolutionary history within prokaryotes before being transferred to eukaryotes (Ponce-Toledo et al., 2017). Testing this hypothesis requires further exploration of the evolutionary links between iACC formation across different bacterial lineages. (ii) iACC-forming bacteria can influence the local geochemical cycling of alkaline earth metals (Ca, Sr, Ba) and other metals, particularly when they bloom as large populations (Blondeau et al., 2018b; Gaetan et al., 2023; Liu et al., 2025). Last, (iii) some of these bacteria are capable of sequestering toxic radioisotopes, opening new avenues for the development of bioremediation strategies (Mehta et al., 2019, 2022a; Pamart et al., 2025).

Despite the significance of this biomineralization process, the molecular mechanisms underlying bacterial iACC formation remain largely unexplored. In *Cyanobacteria*, iACC forms within envelopes thought to consist of proteins and/or lipid monolayers (Blondeau et al., 2018a). Through comparative genomics, a novel gene family named *ccyA* was identified (Benzerara et al., 2022). This gene family lacks homologs with a known function but serves as a diagnostic marker for iACC formation in *Cyanobacteria*. The *ccyA* gene family encodes two-domain proteins, the calcyanins. The C-terminal domain of calcyanins represents a large part of the full-length proteins and is conserved across cyanobacterial clades. It is composed of several long glycine zipper (Gly-zip) motifs. In contrast, the N-terminal domain is much shorter and variable, defining four distinct calcyanin subgroups, corresponding to different *Cyanobacteria* clades (Benzerara et al., 2022). Overall, it is the conserved C-terminal domain that allows to confidently identify members of the calcyanin family despite the variability of their N-terminal domains. A strong correlation has been established between the presence of the C-terminal domain and the capability of a cyanobacterium to form iACC. Interestingly, the N-terminal domain of one calcyanin subgroup contains a CoBaHMA domain, which is hypothesized to play a regulatory role, possibly interacting with anionic lipids (Gaschignard et al., 2024). Furthermore, a recent study suggests that other genes may operate in concert with *ccyA*, such as a *cax* gene encoding a Ca^2+^/H^+^ transporter (Bruley et al., 2025).

In this study, we report the discovery of a homolog of the *ccyA* gene in *Methylococcaceae*, a family of *Gammaproteobacteria*. Furthermore, the two available cultivated strains from this family were found to form iACC, providing evidence that *ccyA* is a marker of iACC biomineralization beyond the phylum *Cyanobacteria*. The phylogenetic distribution of *ccyA* within *Methylococcales*, along with its comparison with the *ccyA* genes in *Cyanobacteria*, offer new insights into the evolutionary origins of iACC biomineralization in *Methylococcaceae*.

## 2. Materials and methods

### 2.1 Genome collection and phylogenomic analyses

The 858 high-quality genome sequences of species of the order *Methyloccoccales* available in the Genome Taxonomy Database (GTDB) version r220 (Parks et al., 2022) were downloaded. They included seven families according to the GTDB taxonomy: *Cycloclasticaceae* (43 genomes), *Methylococcaceae* (112 genomes), *Methylomonadaceae* (679 genomes), *Methylothermaceae* (13 genomes), UBA1147 (6 genomes), UBA2778 (3 genomes), and DRLZ01 (2 genomes). Eight additional gammaproteobacterial genomes were downloaded to serve as outgroup in phylogenomic analyses: *Coxiella burnetii* cb109 (GCA_000300315.1), *Legionella pneumophila* Hextuple_2q (GCA_000277025.1), *Reinekea blandensis* MED297 (GCF_000153185.1), *Dichelobacter nodosus* (GCF_001263335), *Francisella tularensis* FSC147 (GCA_000018925.1), *Nitrosococcus oceani* ATCC 19707 (GCF_000012805.1), *Halorhodospira* sp. (GCA_007121755.1), and *Alkalilimnicola ehrlichii* MLHE-1 (GCF_000014785.1). Protein-coding genes were predicted with Prodigal (Hyatt et al., 2010) when annotations were missing.

GTDB-Tk (Chaumeil et al., 2022) was used to identify 120 conserved protein markers encoded in the whole collection of *Methylococcales* and outgroup genome sequences. Individual markers were aligned using MAFFT L-INS-I v7.310 (Katoh et al., 2019). Poorly aligned sites were removed using trimAl (Capella-Gutiérrez et al., 2009) with the –strictplus option and the resulting trimmed alignments were concatenated using an in-house Python script, yielding an alignment of 36,916 amino acid sites. A Maximum Likelihood tree based on this supermatrix was inferred using IQ-TREE v3.0.1 (Wong et al., 2025) with the LG+G substitution model (Le and Gascuel, 2008). Branch statistical support was calculated with 1,000 ultrafast bootstrap replicates (Hoang et al., 2018). A tree including only the *Methylococcales* species containing the *ccyA* gene was constructed using the same concatenated markers and the same method described above.

### 2.2 Detection of *ccyA*

The detection of *ccyA* was conducted using the in-house tool pCALF (standing for python CALcyanin Finder, https://github.com/K2SOHIGH/pcalf/tree/main), which uses four hidden Markov model (HMM) profiles describing the C-terminal glycine zipper triplication (called the (GlyZip)_3_ motif), specific of calcyanins in *Cyanobacteria* as explained by Gaschignard et al. (2024). These four HMM profiles describe the whole triplication ((GlyZip)_3_), as well as each glycine zipper individually (Gly1, Gly2, and Gly3). They were searched against protein sequences using pyHMMER (Larralde and Zeller, 2023). A sequence is classified as calcyanin when it has significant hits against the GlyX3 HMM profile (sequence coverage threshold: 60%; E-value threshold: 1e^-20^) and against individual Gly1, Gly2, and Gly3 HHM profiles in this specific order (sequence coverage threshold: 70%, E-value threshold: 1e^-10^).

### 2.3 Sequence and three-dimensional (3D) structure analysis

Calcyanin protein sequences detected in *Methylococcales* were compared with those found in *Cyanobacteria*. For this purpose, multiple sequence alignments were performed using MAFFT L-INS-I v7.310 (Katoh et al., 2019), then checked and further analyzed using Hydrophobic Cluster Analysis (HCA) (Bitard-Feildel et al., 2018). Aligned sequences were rendered on the basis of ESPript 3.0 results (Gouet et al., 2003). 3D structure predictions were performed using the AlphaFold3 server (Abramson et al., 2024). Visualization and manipulation of 3D structures were performed using Chimera 1.13.1 (Pettersen et al., 2004). 3D structure similarities were also explored and quantified using TM-align on the RCSB pairwise structure alignment tool (https://www.rcsb.org/alignment; Bittrich et al., 2024).

### 2.4 Bacterial strains

The strain *Methylococcus geothermalis* IM1 (=JCM 33941 =KCTC 72677) was ordered from the Riken bioresource research center in Japan (https://www.jcm.riken.jp/cgi-bin/jcm/jcm_number?JCM=33941. *M. geothermalis* is an aerobic, planktonic and non-motile methanotroph isolated from a hot spring soil in Korea (Awala et al., 2020). Here, it was cultured in triplicates at 37 °C in 10 mL of modified nitrate mineral salts medium-2 (M1221; https://www.jcm.riken.jp/cgi-bin/jcm/jcm_grmd?GRMD=1221) in a 120-mL glass serum bottle with a 25-75 vol% of CH_4_-air mix in the headspace. The genome sequence is available under NCBI accession number CP046565.1.

The strain *Methylococcus mesophilus* 16-5^T^ (=KCTC 82050^T^) was provided by Dr. Samuel Imisi Awala from University of Calgary. *M. mesophilus* is a mesophilic, aerobic methanotroph isolated from a rice field in Korea (Awala et al., 2023). It was cultured in triplicates at 21°C in 10 mL of modified nitrate mineral salts medium-2 (M1221; https://www.jcm.riken.jp/cgi-bin/jcm/jcm_grmd?GRMD=1221) in a 120-mL glass serum bottle with a 25-75 vol% of CH_4_-air mix in the headspace. The genome sequence is available under NCBI accession number CP110921.1.

Cell growth for both strains was assessed by measuring optical density at 600 nm. The pH of the cell suspension was measured over the growth of the cells with a precision better than 0.1.

### 2.5 Fourier Transform infrared spectroscopy (FTIR)

FTIR spectroscopy analyses of both strains were performed to detect bulk signals from iACC inclusions, following the methodology outlined by (Mehta et al., 2022b). Cells were pelleted from 15 mL of a culture grown to an optical density of 0.2, washed with MilliQ® water, dried and ground in an agate mortar. Attenuated total reflectance Fourier transform infrared (ATR-FTIR) spectra were acquired in the mid-IR region (4000–600 cm^-1^) using a NICOLET 6700 FTIR spectrometer equipped with a diamond internal reflection element (IRE), a KBr beam splitter and a deuterated triglycine sulfate (DTGS) detector. Spectra were collected by averaging 64 scans at resolution of 4 cm^-1^.

### 2.6 Synchrotron-based X-ray absorption near-edge structure (XANES) spectroscopy

The relative quantification of Ca in iACC within *M. geothermalis* cells was achieved using XANES spectroscopy, following the approach detailed by Mehta et al. (2023). For this, 15 mL of a cell suspension were harvested by centrifugation, washed, and dried at 37 °C for 48 hours. The dried biomass (∼5 mg) was then diluted in 35 mg cellulose, pressed into a 10 mm diameter pellet, and mounted onto Ca-free holders. XAS at the Ca K-edge was performed on the LUCIA beamline at the SOLEIL synchrotron (Saint-Aubin, France) (Flank et al., 2006; Vantelon et al., 2016). The fixed exit double-crystal monochromator was equipped with Si (111) crystals.

Spectra were recorded in the fluorescence mode using a 60 mm^2^ mono-element silicon drift diode detector (Bruker). In the pre-edge region (3940–4030 eV), spectra were recorded with a step size of 2 eV and a counting time of 1 s. In the edge region (4030–4070 eV), spectra were recorded with a step size of 0.25 eV and a counting time of 1 s. The sample holder was positioned at a 24° angle relative to the fluorescence detector to minimize self-absorption effects and was maintained at a temperature of 77 K using a liquid He-cryostat to minimize potential beam damages and localized heating. Energy calibration was performed using a calcite reference, with the first inflection point set to 4045 eV.

### 2.7 Scanning electron microscopy (SEM)

Cell suspensions of both *M. geothermalis* and *M. mesophilus* were filtered onto 0.22 μm Isopore polycarbonate filters and rinsed with 10 mL of MilliQ® water. The filters were air-dried at ambient temperature and mounted on aluminum stubs using double-sided carbon tape. Before analysis, the filters were coated with a thin layer of carbon via evaporation to make them electrically conductive. Scanning electron microscopy (SEM) analyses were performed using a Zeiss™Ultra55 SEM with an electron accelerating voltage of 10 kV, a working distance of 7.5 mm and a 60 μm aperture. Images were captured using a backscattered electron detector (AsB detector). Energy-dispersive X-ray spectrometry (EDXS) data were acquired with a Bruker QUANTAX EDS detector, enabling hyperspectral imaging (HyperMap), and the data were analysed with the Esprit software (Brucker).

### 2.8 Transmission electron microscopy

A 0.5 mL cell suspension was harvested by centrifugation, rinsed twice in 1 mL of MilliQ® water before resuspension in 0.5 mL of MilliQ® water. A drop of 3 μL was deposited onto a formvar-coated TEM grid and air-dried. Observations were carried out using a Jeol 2100F TEM microscope operating at 200 kV, equipped with a Schottky emitter, a STEM device for Z-contrast imaging in high-angular annular dark-field (HAADF) mode, and a Jeol Si(Li) X-ray detector for elemental analyses via energy dispersive x-ray spectrometry (EDXS).

### 2.9 Synchrotron-based cryo-soft X-ray tomography (c-SXT)

Three-dimensional X-ray imaging of *M. geothermalis* cells was performed at the ALBA synchrotron (Barcelona, Spain) using cryo-transmission X-ray microscopy at the MISTRAL beamline (Sorrentino et al., 2015). The cryofixation process involved plunge-freezing at the Institut Pasteur (Paris, France) using a Leica EM GP. A Quantifoil TEM grid (Q-R2/1-C2100) was loaded in the chamber of the plunge freezer. Four microliters of a two-day-old *M. geothermalis* culture (OD = 0.251) and one microliter of a 100-nm gold nanoparticles suspension (BBI Solutions, 5X concentrated) were deposited on the front of the grid. After blotting for four seconds from the back, grids were plunged into a container filled with liquid ethane cooled with liquid nitrogen (−196°C), ensuring proper vitrification of the samples. Subsequently, the samples were shipped from Paris to the ALBA synchrotron in a dry shipper and transferred to the cryo-chamber of the Mistral beamline for imaging.

A tilt series of projections was collected from −70° to +70° in 1° increments (except otherwise mentioned), with an incident X-ray energy of 520 eV. Each projection was exposed for 3 seconds. A 40-nm Fresnel zone plate was used, achieving an effective pixel size of 12 nm. The projections were normalized for incoming flux and deconvolved using the measured point spread function of the optical system (Otón et al., 2016). Projections were aligned using Etomo with the 100-nm gold nanoparticle fiducials. Tomographic reconstruction, along with simultaneous iterations reconstruction technique (SIRT) deconvolution, were performed using IMOD (Otón et al., 2016). Volume segmentation, calculations and visualization of tomograms were carried out using the Microscopy Image Browser and ImageJ (Belevich et al., 2016).

## 3. Results

### 3.1 Detection and phylogenetic distribution of *ccyA* in the order *Methylococcales*

The *ccyA* gene (more precisely, the long C-terminal domain of the encoded calcyanin protein) has been identified as a diagnostic marker for iACC biomineralization in *Cyanobacteria*. Moreover, homologs of *ccyA*, identified based on the sequence of the calcyanin C-terminal domain, were also reported in four gammaproteobacteria of the order *Methylococcales* in a previous study (Benzerara et al., 2022)). However, the evolutionary relationship between the *ccyA* genes of cyanobacteria and those of *Methylococcales* remains unclear. Furthermore, it is unknown whether *Methylococcales* possessing *ccyA* are capable of forming iACC, as cyanobacteria do. To investigate these questions further, we systematically searched for *ccyA* homologs across the 858 genome sequences of *Methylococcales* species available in the Genome Taxonomy Database (GTDB) version r220 (Parks et al., 2022) (Table S1). This search was performed using the pCALF tool (see Materials and methods) with the E-value, coverage and identity thresholds provided in Table S2.

We detected *ccyA* in 39 genomes of species belonging to the order *Methylococcales* (out of 858 available genomes, see Table S1), distributed in the genera *Methylococcus* (including *M. geothermalis* and *M. mesophilus*), *Methylospira* (*Candidatus* Methylospira mobilis), and *Methylumidiphilus* (many strains affiliated with *M. alinensis*) (Table 1). According to the GTDB taxonomy, all these species were assigned to the family *Methylococcaceae* (represented by a total of 112 genomes; *ccyA-*containing genomes thus represent 35% of the available *Methylococcaceae* genomes); none were associated with the species-rich *Methylomonadaceae* (679 genomes) or with the other families of the order *Methylococcales* (Figure 1 and Figure S1). From the observed phylogenetic distribution of *ccyA* in *Methylococcaceae*, the most parsimonious scenario thus suggests that *ccyA* appeared in a deep branch of *Methylococcaceae* after the divergence from *Methylomonadaceae* (Figure 1). The species possessing *ccyA* occur in diverse environments, including the water column and sediment of freshwater lakes, compost, active municipal landfills, and rice field soils (Table 1 and Table S2). In all identified genomes, the protein calcyanin was consistently annotated as a hypothetical protein. According to the GTDB metadata, most of the genomes containing *ccyA* (34 out of 39) were metagenome-assembled genomes (MAGs), and only five originated from cultured isolates (Table 1). Only two of these isolates were available: *Methylococcus geothermalis* IM1 and *Methylococcus mesophilus* 16-5, which we further studied to search for their capability to biomineralize iACC.

**Figure 1:**
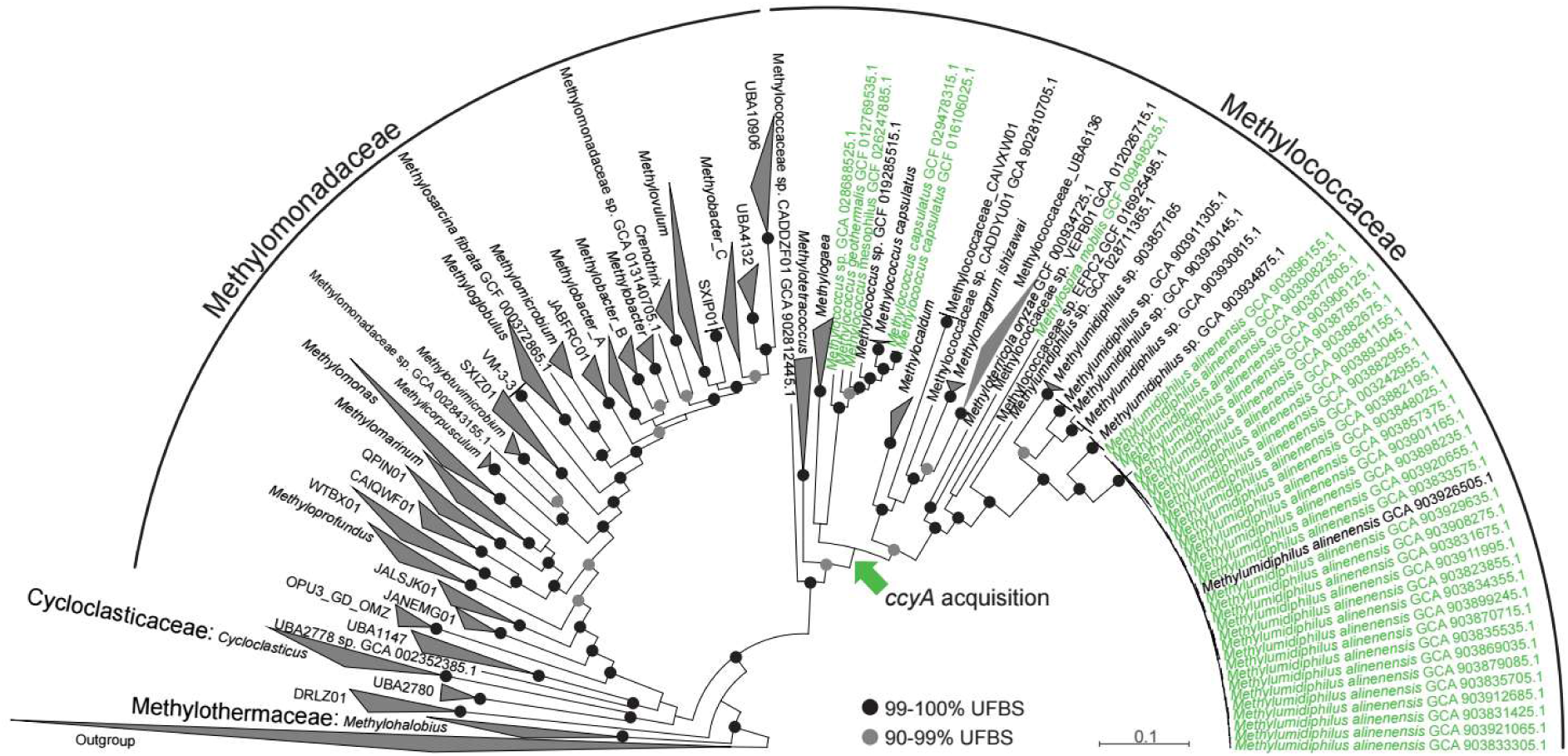
Maximum likelihood phylogenomic tree of 858 *Methylobacterales* taxa available in the Genome Taxonomy Database (GTDB version 220) (see Table S1). The tree is based on the concatenation of 120 conserved proteins. Taxa containing the *ccyA* gene are highlighted in green. Taxa that do not contain the *ccyA* gene are collapsed. Taxon names correspond to the GTDB taxonomy. The green arrow indicates the most likely point (based on maximum parsimony) for the acquisition of *ccyA* by an ancestor of several *Methylococcaceae* lineages (earlier acquisitions would imply an increasing number of subsequent independent losses). Ultrafast bootstrap support (UFBS) values >90% are indicated on branches (black dots: 99-100%; gray dots: 90-99%). The complete, un-collapsed tree is available in Fig. S1.

**Table 1:**
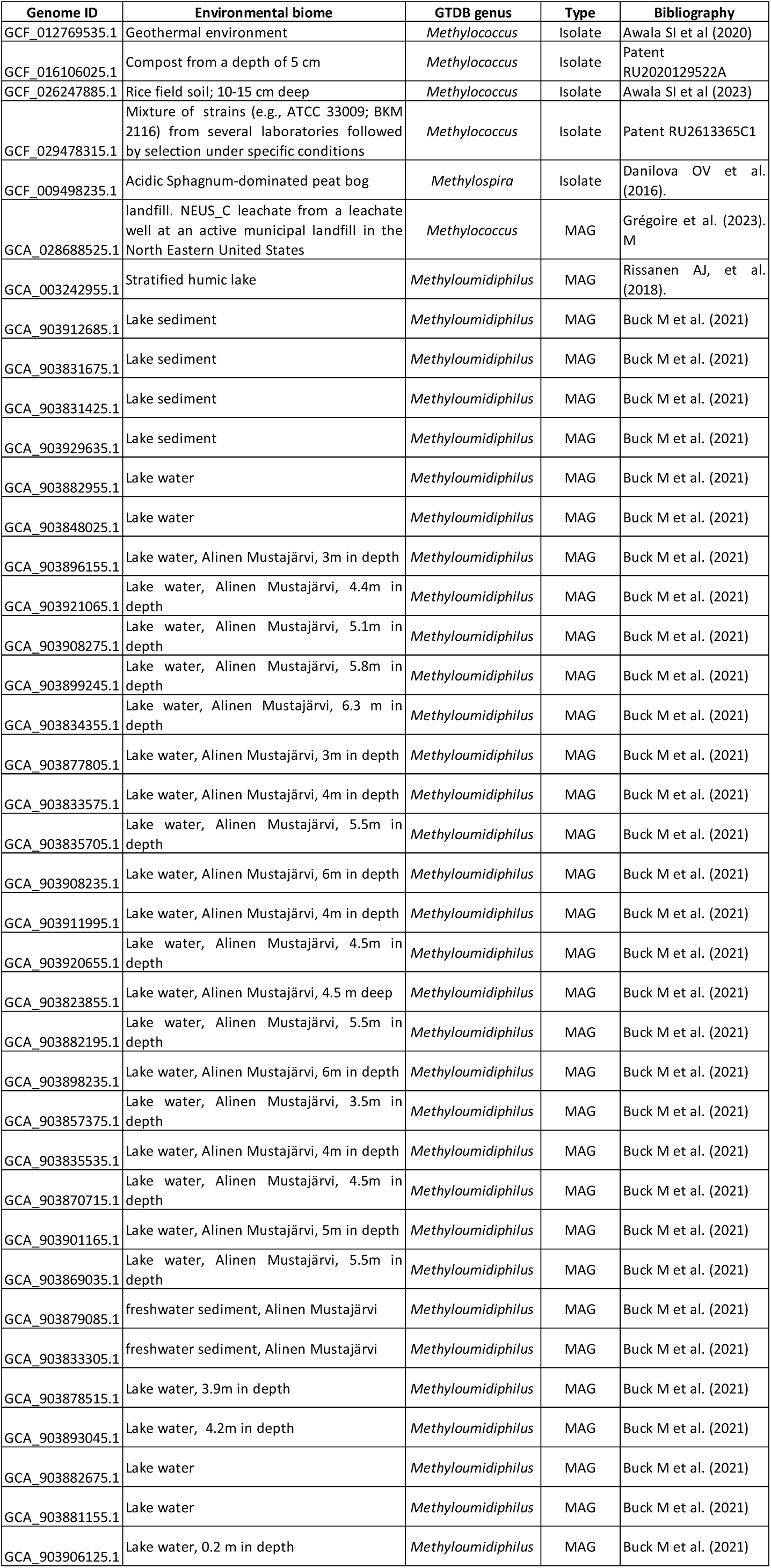
List of genomes of *Methylococcales* containing *ccyA*. Main information about the genomes, including whether they are derived from cultivated strain (isolate) or metagenomes (MAG), the genera they are affiliated with, following GTDB classification, and their environmental origin. Additional information is provided in Table S2.

### 3.2 Comparison of *Cyanobacteria* and *Methylococcaceae* calcyanin sequences

The calcyanin protein comprises two distinct domains: the N-terminal domain is short and variable, and defines four calcyanin types in Cyanobacteria, called W (or CoBaHMA), X, Y and Z, with the latter three restricted to specific clades (*Gloeomargarita, Fischerella* and closely related genera, and *Microcystis*, respectively). In contrast, the C-terminal (GlyZip)_3_ domain is much longer, characterized by a triplicated long glycine zipper, and it is highly conserved across cyanobacteria. The conservation of this domain allows for the unambiguous identification of calcyanin based solely on its sequence (Benzerara et al., 2022; Gaschignard et al., 2024).

A multiple sequence alignment of the C-terminal domains revealed conservation of the signature motifs of the (GlyZip)_3_ domain in *Methylococcaceae* (Figure 2). Key features included the repeated basic Gly-X(3)-Gly pattern within each GlyZip motif, interrupted at the midpoint by a highly conserved Gly-Pro dipeptide. Additionally, several polar and aromatic amino acids were notably conserved, particularly in the third GlyZip motif (indicated by stars in Figure 2). Some variability was observed among *Methylococcaceae* members, principally occurring outside the GlyZip motifs, similar to the variability observed across cyanobacteria. Sequence comparisons further revealed that the *Methylococcaceae* C-terminal (GlyZip)_3_ domains were more similar to Z-type calcyanins of *Microcystis* (47 % identity) than to other calcyanin types found in other cyanobacteria (30 % identity). High and specific sequence similarities were especially apparent in the first GlyZip motif of *Methylococcaceae* and *Microcystis* (∼60 % sequence identity versus lower than 40 % for other Cyanobacteria, Figure 2). Additionally, the N-terminal domains of *Microcystis* Z-type calcyanins and *Methylococcaceae* calcyanins also shared several features: they lacked strongly hydrophobic amino acids and were enriched in alanine and polar residues (Figure S2). This property is also shared by the N-terminal sequences of type-X and type-Y calcyanins. However, despite these compositional similarities, no robust sequence alignments could be established between the *Microcystis* and *Methylococcaceae* N-terminal sequences, as it was also the case between *Methylococcaceae* and type X and type Y N-terminal sequences. We note that similarly, no significant sequence alignments could be established between the N-terminal sequences of the different cyanobacterial calcyanins, the so-called CoBaHMA-, X-, Y- and Z-type calcyanins and yet they are all markers of iACC biomineralization (Benzerara et al., 2022).

**Figure 2:**
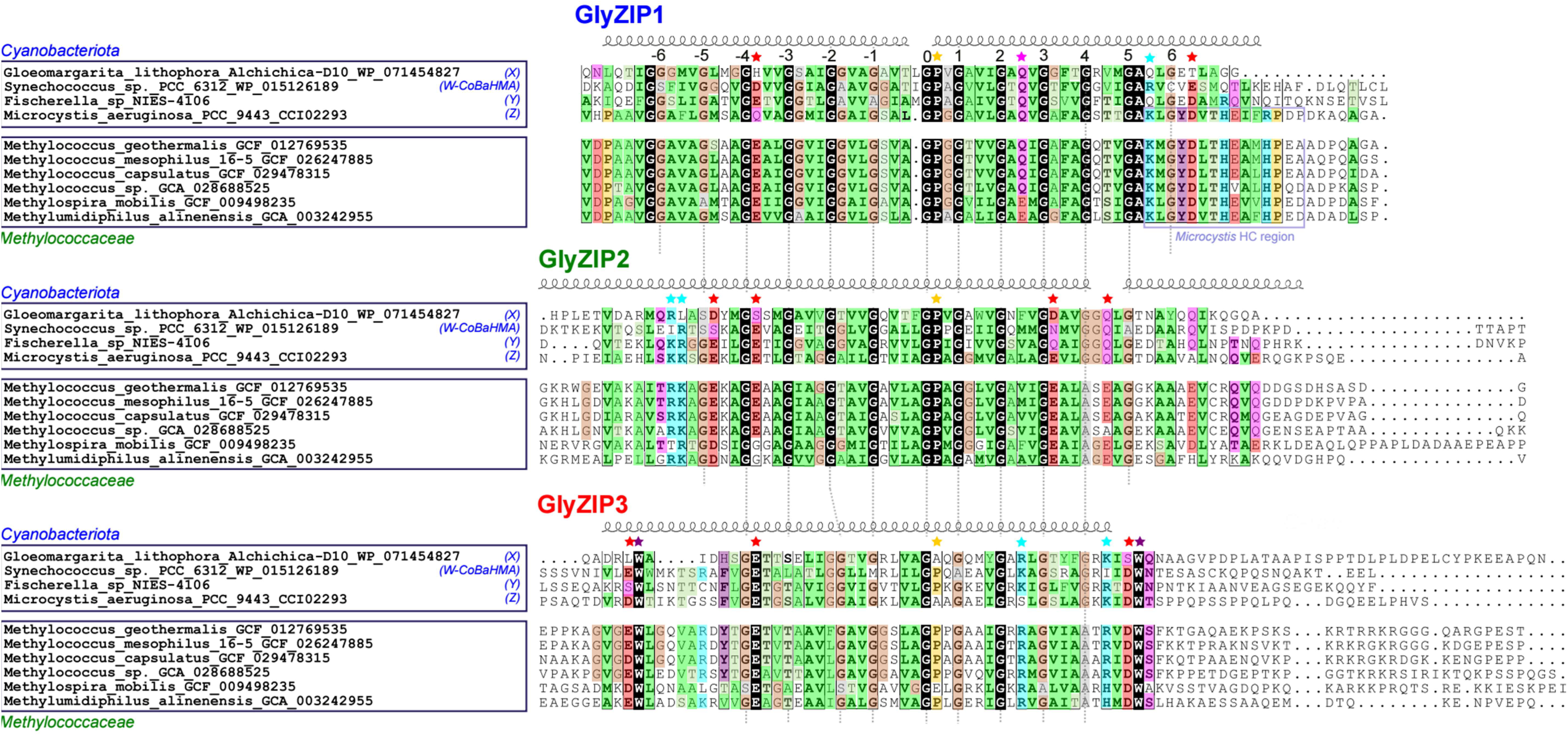
Comparison of the sequences of calcyanin GlyZip motifs of *Methylococcaceae* and cyanobacteria. Four calcyanin sequences representative of the different clades were selected for cyanobacteria, while for *Methylococcaceae,* six non-redundant ones were extracted from Table 1. The *Fischerella* sp. NIES-4106 calcyanin sequence was chosen as it has three GlyZip motifs, instead of two for most of the calcyanin sequences of this clade. Identities are shown in white on a black background, while similarities are highlighted with colors, according to amino acid properties and position in the alignment (with highly similar positions indicated in bold). The periodic motif involving glycine (or a small amino acid) every four residues is highlighted by numbers relative to the central conserved proline (gold star), and can be followed from one GlyZip motif to the next using vertical dotted lines. Highly conserved polar/aromatic positions are highlighted with stars. The lilac pink box highlights a region which is highly conserved (HC) between the *Microcystis* and *Methylococcaceae* sequences. The secondary structures (α-helices) as predicted by AlphaFold3 (pLDDT<70) on the *Methylococcus geothermalis* sequence are shown on top of the alignment.

Three-dimensional (3D) structure predictions generated by AlphaFold3 suggested that the three GlyZip motifs comprising the C-terminal domain of calcyanin share a common hairpin structure (Figure 3). This structure consists of two tightly packed alpha-helices, whose close packing is provided by glycine zippers (Figure 3). Even though the 3D structure of the GlyZip motifs are predicted with low pLDDT values (<50), confidence can be placed in the models of the GlyZip core region, spanning positions −4 to 4, over approximately 40 residues. This core region exhibits structural similarity across the three GlyZip motifs within a single species and among different species (see the RMSD values in Table S3). In contrast, structural variability is observed in the AlphaFold3 models, even those built for a same sequence, particularly at the N- and C-terminal extremities (thus outside the GlyZip core). The sole exception is GlyZip1, for which the glycine zipper and associated helix packing invariably extends to these terminal regions across all models (Figure 3). Consequently, only the entire GlyZip1 motif permits structural comparisons between species, given the stability of its 3D structure predictions across its full length. In contrast, the variability observed in models of a same sequence precludes any meaningful comparison of the GlyZip2 and GlyZip3 motifs. These analyses revealed that the 3D structure of GlyZip1 in *Methylococcaceae* is structurally more similar to that of *Microcystis* than to those of other cyanobacteria (Figure 3 and metrics in Table S3). This similarity extends to the N-terminal domains, located upstream of the (GlyZip)3 domains, which are predicted to form alpha-helices structures, though with low confidence scores (Figure S2) and without any possibility to align the sequences. Importantly, AlphaFold3 failed to accurately predict the spatial arrangement between the three GlyZip motifs, indicating that the (GlyZip)_3_ domain may adopt an as-yet-unknown 3D fold.

**Figure 3:**
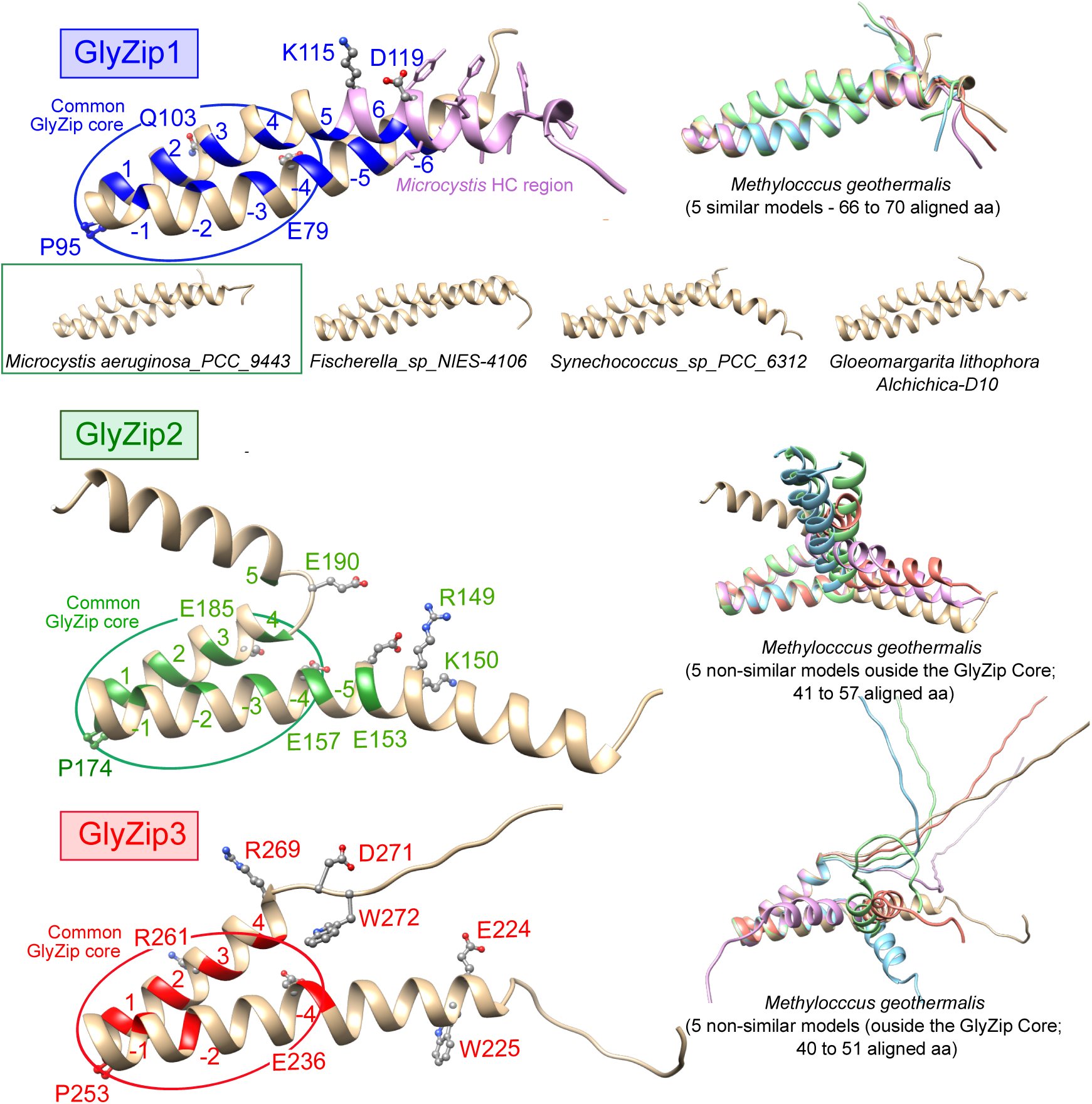
Ribbon representation of the 3D structure of the GlyZip motifs, as predicted by Alphafold3 using the *Methylococcus geothermalis* calcyanin sequence. The GlyZip motifs were superimposed for comparative analysis (metrics in Table S3). The periodic motifs involving glycine (or a small amino acid) every four residues are colored on the ribbons. Highly conserved charged/polar/aromatic residues (stars in Figure 2) are depicted in atomic details. The highly conserved (HC) region between the *Microcystis* and *Methylococcaceae* sequences (Figure 2) is highlighted in lilac pink. For note, the Alphafold3 per-residue pLDDT values are < 50, probably due to the fact that no template could be found in the Protein Data Bank to support the relevance of this prediction (and not because of a dubious prediction, a hypothesis ruled out by in-depth analysis of the model, in particular highlighting the common central GlyZip core around the Gly-Pro sequence). Structural uncertainties concern the N- and C-terminal extremities of the GlyZip motifs 2 and 3, which considerably vary (even in the five models built based on the same sequence from *Methylococcus geothermalis* – shown at right, mean RMSD and TM scores of 2.88 Å / 0.57 (41 to 57 aligned aa) for GlyZip2 and 2.57 Å / 0.43 (40 to 51 aligned aa) for GlyZip3), and which deviate from the packing imposed by the periodic repetitions of glycine. In contrast, GlyZip1 motif 3D structures are similar across the models of *M. geothermalis* calcyanin (1.57 Å / 0.86 (66 to 70 aligned aa)) and across the calcyanins of different species, as supported by RMSD values and TM-scores reported in Table S3. Overall, the GlyZip1 motifs and the GlyZip cores of GlyZip2 and GlyZip3 motifs in *Methylococcus geothermalis* exhibit greater structural similarity to those of *Microcystis aeruginosa* calcyanin than to calcyanins from other cyanobacteria.

Moreover, the similarity between the Z-type calcyanins of *Microcystis* and the newly identified calcyanins of *Methylococcaceae* is strongly supported by phylogenetic analysis. Indeed, a maximum likelihood tree constructed using the conserved GlyZip motifs in the C-terminal domain of all calcyanins, showed that the *Microcystis* and *Methylococcaceae* sequences are closely related with maximal statistical support (Figure 4). Interestingly, the phylogenetic relationships among the *Methylococcaceae* GlyZip sequences perfectly mirrored those derived from the phylogenomic analysis based on a large set of conserved markers from the corresponding *Methylococcaceae* species (Figure 4). This mirroring indicates that the *ccyA* gene has followed the speciation history of these species.

**Figure 4:**
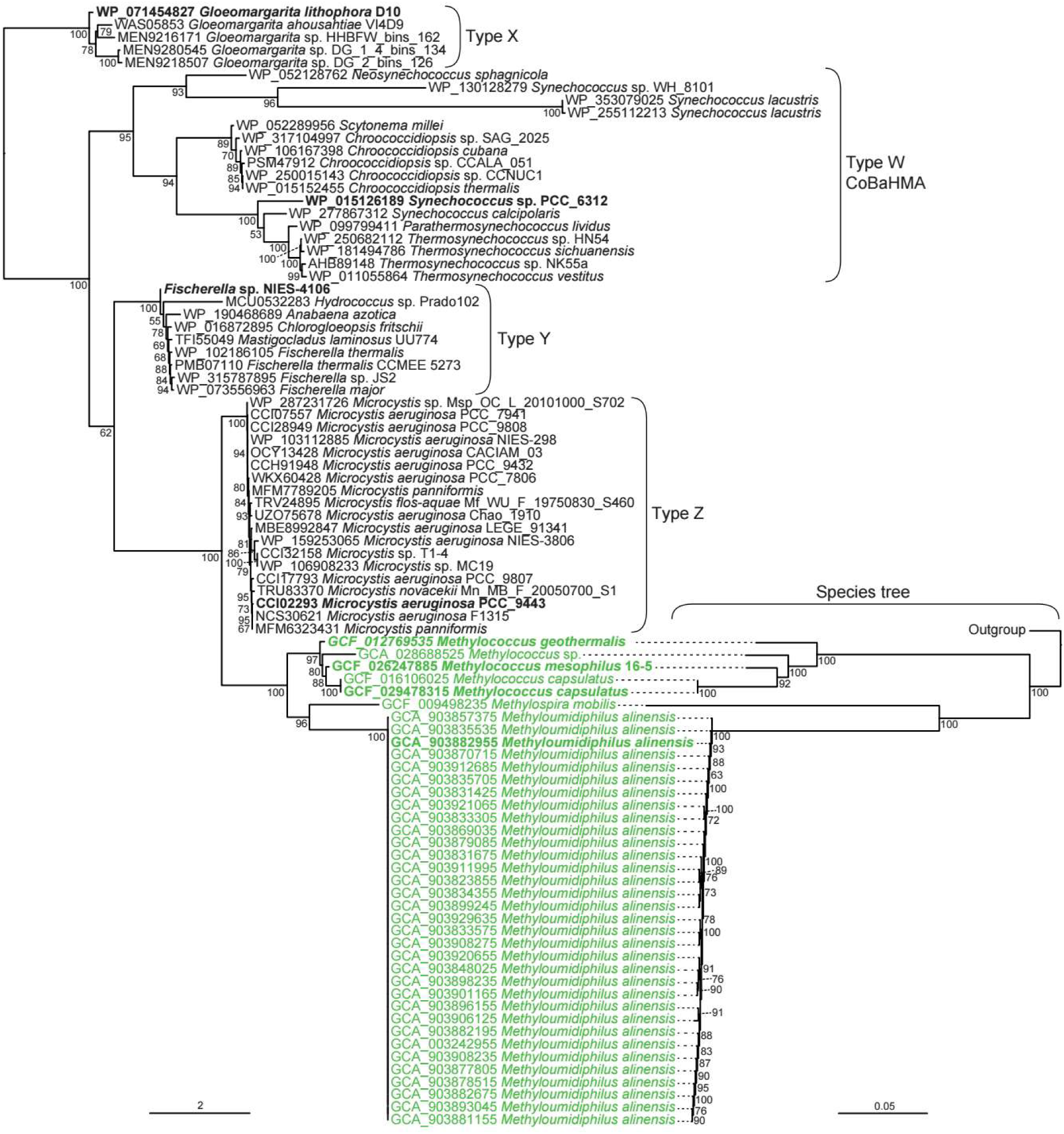
Maximum likelihood phylogenetic tree of the GlyZip domain of the calcyanin sequences. Calcyanin sequences of *Methylococcaceae* species are highlighted in green. The different types of cyanobacterial calcyanins (X, W/CoBaHMA, Y, and Z) are indicated. For the *Methylococcaceae*, a species tree based on the concatenation of 120 conserved markers is shown on the left (this tree is rooted on an outgroup composed of *Methylococcaceae* sp. GCA_902812445.1, *Methylotetracoccus* sp. GCA_025458755.1, *Methylotetracoccus* sp. GCA_004168015.1, and *Methylotetracoccus oryzae* GCF 006175985.1). Numbers on branches show ultrafast bootstrap values. The sequences used in the comparison of calcyanin sequences in Figure 2 appear in bold here.

### 3.3 Experimental evidence of iACC formation by *Methylococcaceae* cultivated strains

Both genomes of the cultivated strains *Methylococcus geothermalis* IM1 and *Methylococcus mesophilus* 16-5 contain the *ccyA* gene (based on the signature of the C-terminal domain of calcyanin), which is indicative of iACC formation in cyanobacteria. Consequently, we searched for the presence of iACC in these strains. Laboratory cultures exhibited a typical exponential growth curve, with a doubling time of 5.8±0.1 h for *M. geothermalis* and 13.1±0.8 h for *M. mesophilus* (Figure S3).

Attenuated total reflectance Fourier transform infrared (ATR-FTIR) spectroscopy performed on cell pellets showed unambiguously the presence of ACC in both *Methylococcaceae* cultivated strains (Figure 5 and Table S4 for band assignment). Indeed, spectra showed a band at around 860 cm^-1^, detected in both *M. geothermalis*, *M. mesophilus*, a synthetic ACC reference and iACC-forming cyanobacteria, but absent in a reference calcite and non-iACC-forming cyanobacteria, and that was shown to be characteristic of ACC (Mehta et al., 2022b). Other features observed in the spectra were bands at 1641 and 1528 cm^-1^, identified as amide I and amide II bands (Figure 5). Bands at 1444 and 1391 cm^-1^ detected in both *M. geothermalis* and *M. mesophilus*, are similar to those observed at 1474 and 1395 cm⁻¹ in ACC but they are not exclusive to iACC-forming cyanobacteria and likely correspond to methyl groups in lipids and/or C-O groups in carboxylic acids.

**Figure 5:**
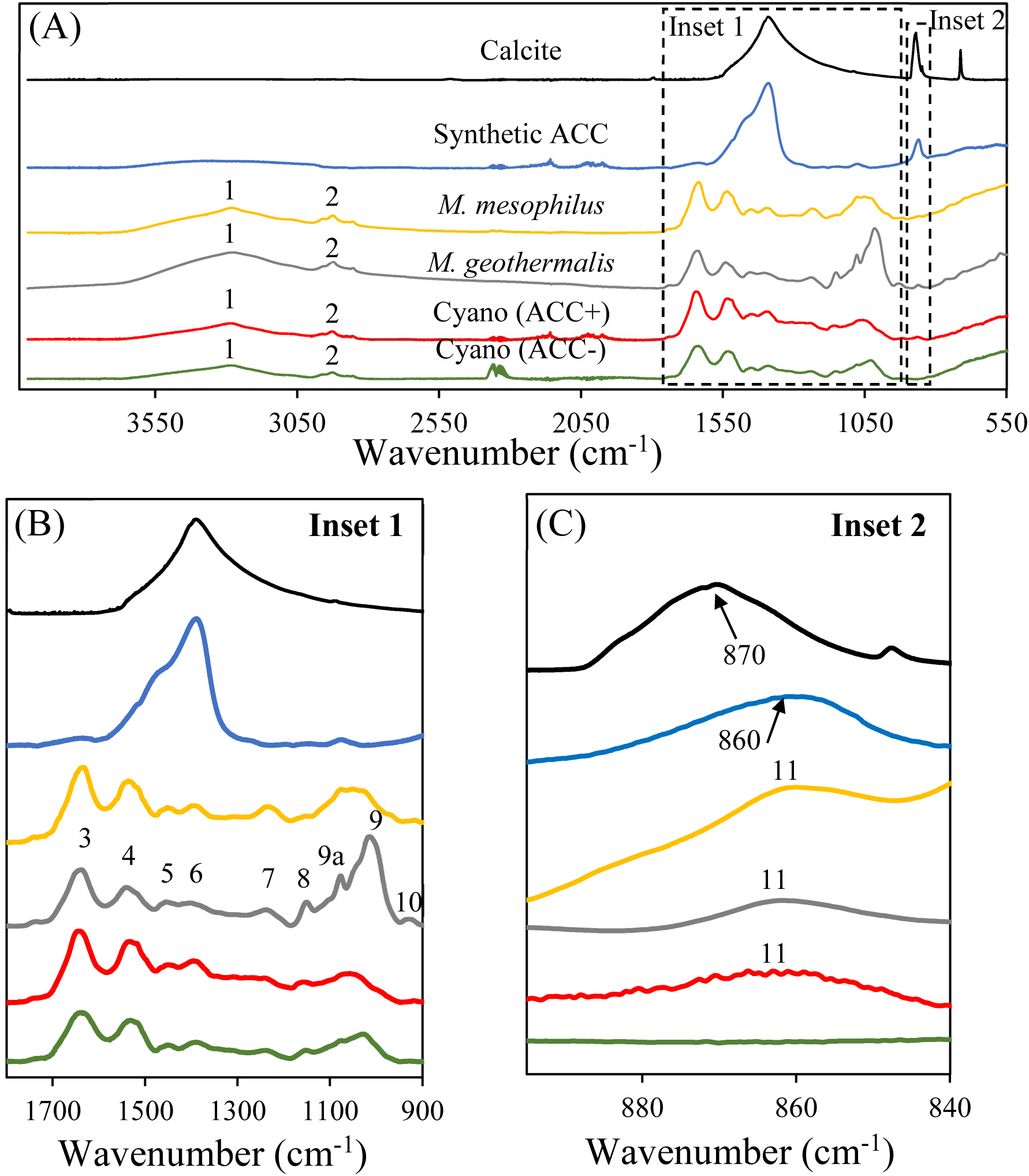
(A) ATR-FTIR spectra of *Methylococcus geothermalis* and *M. mesophilus* in the mid-IR range compared with reference spectra. Reference spectra are given for calcite, synthetic ACC, iACC-forming cyanobacteria (Cyano(ACC+)) and non-iACC-forming cyanobacteria (Cyano(ACC-)). The dashed rectangle envelops the zoomed regions of mid-IR ACC spectra shown in the inset 1 (1800–900 cm^-1^) in (B), and inset 2 (950–820 cm^-1^) in (C). Numbers on the spectra correspond to the band numbers and their assignments are provided in Table S4. Band 11 at 860 cm^-^1 is indicative of ACC.

This finding was further corroborated by X-ray absorption near-edge structure (XANES) spectroscopy at the Ca K-edge conducted on *M. geothermalis* cells. The derivative of the Ca K-edge XANES spectrum of *M. geothermalis* cell pellets closely matched that of synthetic ACC and iACC in cyanobacteria, distinguishing it from calcite and non-iACC-forming cyanobacteria (Figure 6; Mehta et al., 2023). Specifically, the B1 feature in the derivative spectrum, which was shifted compared to calcite, indicated the presence of ACC. Linear combination fitting (LCF) of the XANES data revealed that approximately ∼57% of the Ca in *M. geothermalis* was in the form of ACC, with the remaining 43% of Ca likely complexed by organics and/or associated with polyphosphates (Figure 6).

**Figure 6:**
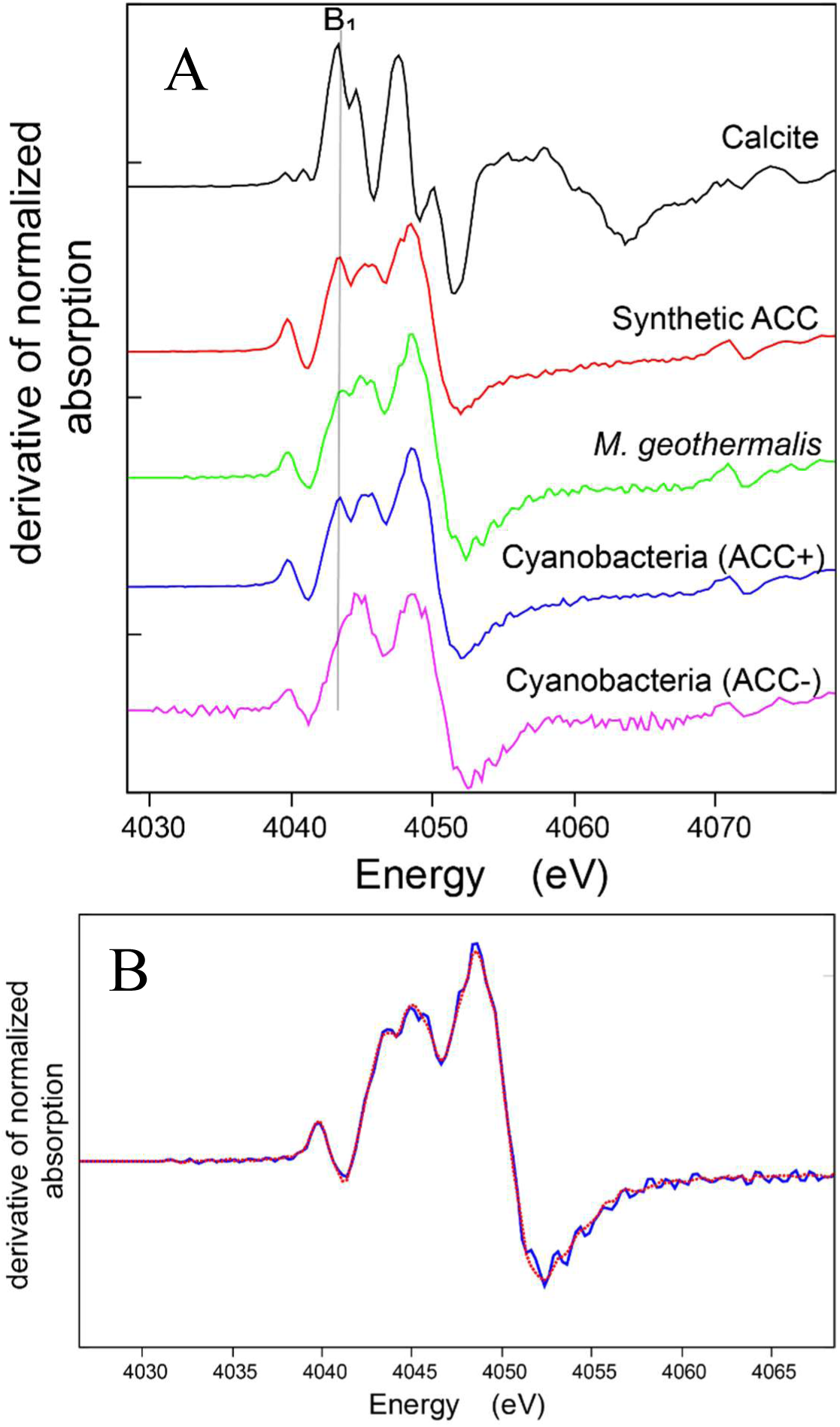
(A) Derivate XANES spectra at the Ca K-edge of *Methylococcus geothermalis* cell pellets compared with reference derivate spectra of calcite, synthetic ACC, iACC-forming cyanobacteria and non-iACC-forming cyanobacteria. The presence of a peak noted B1 indicated by a dashed line is indicative of ACC. (B) Linear combination fitting (LCF) of the derivate XANES spectrum of *Methylococcus geothermalis* cell pellets at the Ca K-edge. The solid line shows experimental data and the dotted line corresponds to the LCF result.

Consistently, both scanning electron microscopy (SEM) and scanning transmission electron microscopy (STEM) in the high-angle annular dark-field (HAADF) imaging mode combined with energy dispersive x-ray spectrometry (EDXS) showed that cells of both strains contained Ca-rich inclusions with some Mg but no S and no P, interpreted as calcium carbonate inclusions (Figure 7 and Figure S4-6). Besides, cells also contained granules rich in phosphorus (P), with some Mg and K, interpreted as polyphosphate inclusions (Rivas-Lamelo et al., 2017).

**Figure 7:**
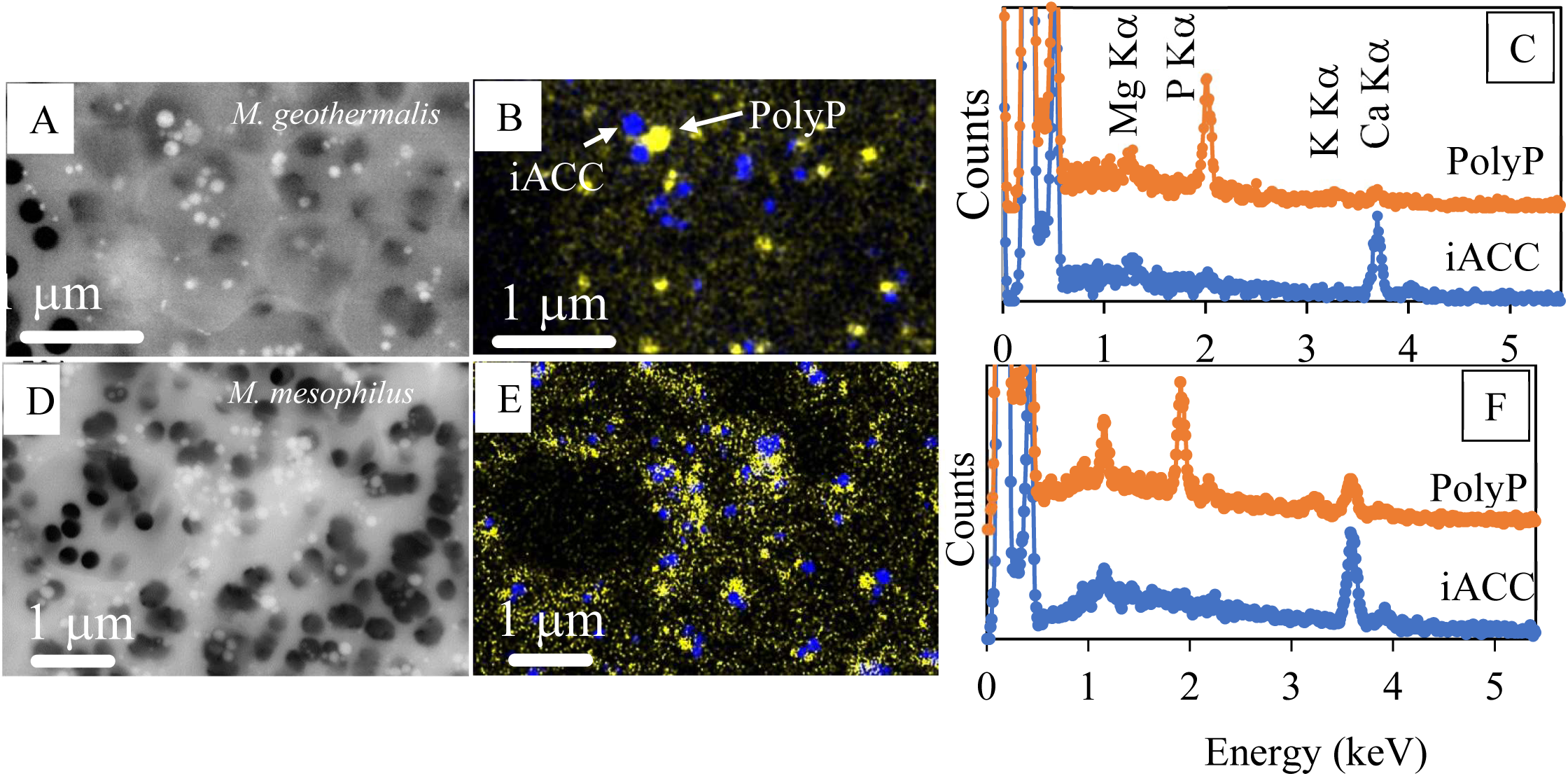
Scanning electron microscopy analyses of cultivated *Methylococcaceae*. (A-C) *Methylococcus geothermalis*. (A) Backscattered electron image. (B) Overlay of Ca (blue) and P (yellow) EDXS maps showing iACC and polyphosphate inclusions, respectively. (C) Characteristic EDXS spectra of polyphosphate (orange) and iACC (blue) inclusions. Emission lines of Mg Ka, P Ka, K Ka and Ca Ka are indicated. (D-F) Same analyses for *M. mesophilus*. More complete datasets can be viewed in the Supplementary Figures.

Last, cryo-soft X-ray tomography (c-SXT) was used to image vitrified *M. geothermalis* cells in a near-native state (15 cells in total, 6 cells used for volume reconstruction). Ca-rich inclusions were observed as regions of higher intensity (Figure 8A and Figure S7). These inclusions are most evident in the tomograms due to their strong absorption contrast and small, spherical morphology (Figure 8A). The absorption contrast from Ca-rich inclusions is much higher than other structures inside the cell (Figure 8B). Clearly defined Ca-rich inclusions were segmented to determine their average size and distribution, yielding a mean diameter of 118 ± 45 nm (n = 58) (Figure 8C). Other structural features, such as intracytoplasmic membranes and phosphate inclusions (Awala et al., 2020), can be distinguished from the smaller Ca-rich inclusions in the tomograms but not from each other due to their similar linear absorption contrast coefficients (beige structures in Figure 8D). For whole-cell segmentation, these structures were therefore grouped as a single material. Figure 8D shows the 3D volume reconstruction of the two cells from Figure 8A, highlighting the outer cell membrane, intracytoplasmic membranes/phosphate inclusions, and Ca-rich inclusions. At least 10 Ca-rich inclusions were found per cell.

**Figure 8:**
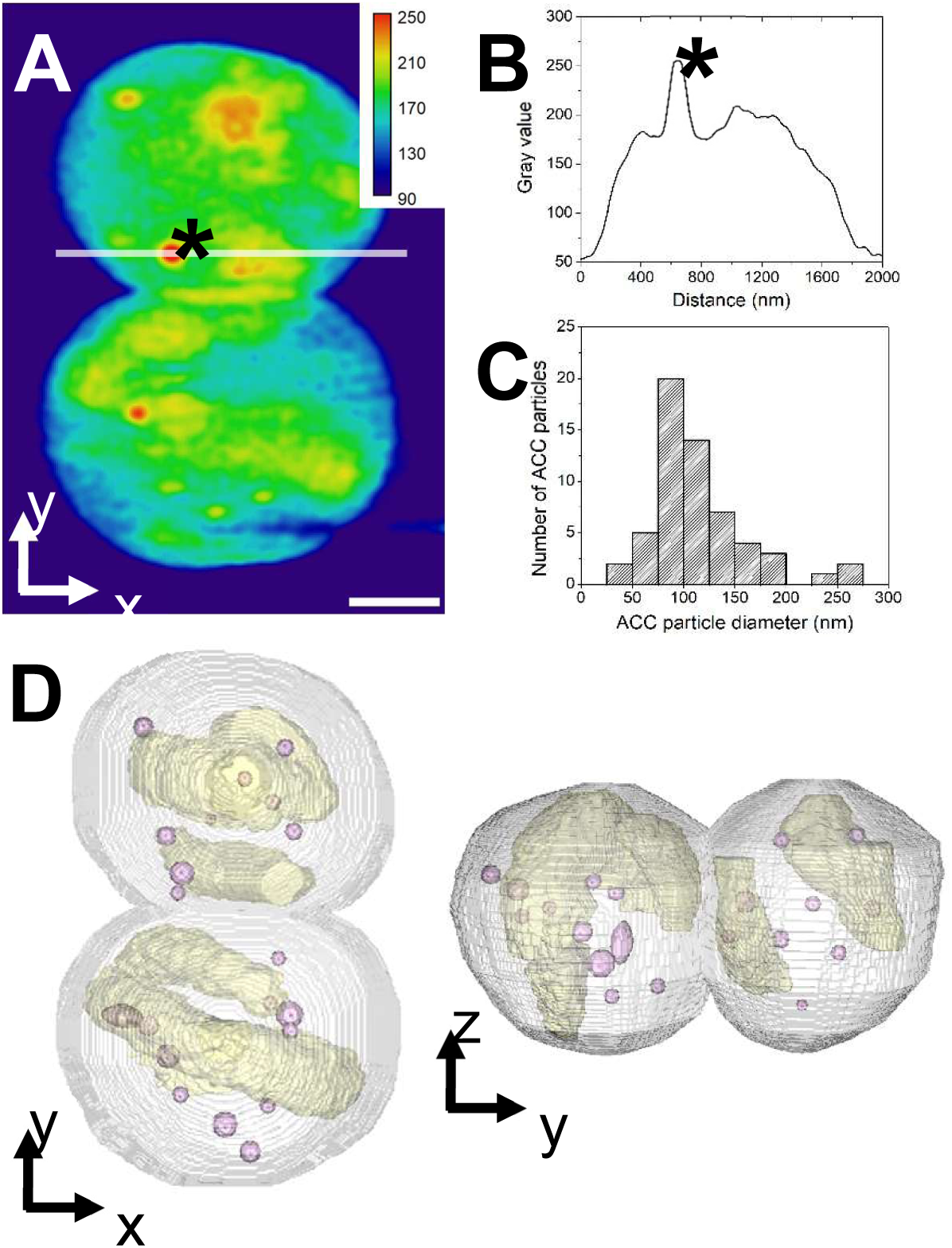
Cryo-soft X-ray tomography (c-SXT) of two ice-vitrified *Methylococcus geothermalis* cells. The image intensity corresponds to X-ray absorption by intracellular material at 520 eV—an energy within the “water window,” where absorption by C and N atoms provides contrast of biological structures without chemical staining. At this energy, absorption by O atoms is minimal, allowing X-ray transmission through both the ice layer and hydrated cells. In contrast, the Ca L₂,₃-edge at 350 eV enables detection of Ca atoms within the cells. (A) Virtual mid-cell z-slice of two *M. geothermalis* cells in reconstructed tomogram (an additional tomogram of two different cells is shown in Figure S3, along with tomogram videos for each dataset). Intensity scale (rainbow colour code) refers to absorption values. Asterisk marks the position of a Ca-rich inclusion. Scale bar is 500 nm. (B) Absorption profile extracted from horizontal line in (A) (asterisk marks the position of a Ca-rich inclusion). (C) Particle size distribution of Ca-rich granules measured from segmented volumes of the tomogram. (D) Volume reconstruction of cells in (A) shown in two perspectives with Ca-rich granules in pink, intracytoplasmic membranes/phosphate inclusions in beige and outer cell membrane in grey.

## 4. Discussion

### 4.1 Formation of iACC inclusions by bacteria remains an overlooked process

Here we report the discovery of iACC in members of the *Methylococcaceae*. This clade expands the growing list of bacteria known to biomineralize ACC, which already includes *Achromatium* (Benzerara et al., 2021; Gray, 2006; Schewiakoff, 1893), numerous phylogenetically diverse members of the cyanobacteria (Benzerara et al., 2022), several magnetotactic *Pseudomonadota*, spanning both *Gammaproteobacteria* and *Alphaproteobacteria*, and magnetotactic *Nitrospirota* (Liu et al., 2021, 2025; Mangin et al., 2025; Monteil et al., 2020). Collectively, these discoveries suggest that iACC biomineralization is more widespread than previously recognized but has often gone unnoticed, likely due to the analytical challenges associated with detecting ACC. Indeed, amorphous carbonates are highly labile and require dedicated techniques for reliable identification (Addadi et al., 2003).

For example, although the ultrastructures of *M. geothermalis* and *M. mesophilus* had previously been examined using electron microscopy, iACC granules were not reported (Awala et al., 2020, 2023). This omission can be explained both by the distinct objectives of those studies that did not target mineralogy and the absence of chemical analyses suited to the detection of iACC. In contrast, our combined use of STEM-EDXS on intact, untreated cells together with Ca K-edge XANES and/or FTIR spectroscopy enabled the unambiguous detection of iACC in both species. Moreover, iACC granules were likely lost during the ultramicrotomy preparations employed by Awala et al. (2020 & 2023) since standard protocols involving chemical fixation, post-fixation and dehydration are known to dissolve iACC in cyanobacteria (Blondeau et al., 2018a; Li et al., 2016). It is therefore possible that the numerous electron-translucent vesicles or intracellular inclusions previously interpreted as polyphosphate or glycogen granules by Awala et al. (2020, 2023) actually represent dissolved or transformed iACC granules. However, this hypothesis can only be rigorously tested using cryo-electron microscopy of vitreous sections (CEMOVIS).

An additional and pivotal line of evidence for iACC biomineralization in the *Methylococcaceae* was the detection of the *ccyA* gene (and more specifically the portion encoding the characteristic C-terminal (GlyZip)_3_ domain of calcyanin), previously identified as a diagnostic marker of iACC formation in cyanobacteria (Benzerara et al., 2022). Our findings extend the diagnostic value of *ccyA* beyond cyanobacteria, although so far, no other clades represented in current genomic databases appear to carry this gene—unless their versions of *ccyA* have diverged substantially. Importantly, however, the absence of *ccyA* in other bacterial lineages does not necessarily imply the absence of iACC formation. For instance, some magnetotactic *Pseudomonadota* and *Achromatium* both form iACC despite lacking *ccyA*, suggesting that multiple, distinct molecular pathways may give rise to iACC biomineralization (Mangin et al., 2025).

Last, numerous MAGs of iACC-forming *Methylococcaceae* that we have detected originate from chemically stratified boreal lakes and ponds (Table 1; Buck et al., 2021). Interestingly, MAGs from other bacteria phylogenetically close to iACC-forming magnetotactic bacteria were also found in the same stratified lakes and ponds in Finland and Sweden (Mangin et al., 2025), suggesting that iACC biomineralization might be a geochemically important process in these lakes.

### 4.2 Relationship between *Methylococcaceae* metabolism and iACC biomineralization

The mechanisms governing iACC biomineralization remain poorly understood. It has been proposed that bacterial primary metabolism may play an important role in driving this process (Görgen et al., 2020). In *Achromatium*, a sulfur-oxidizing bacterium, iACC formation is linked to sulfide oxidation to elemental sulfur (Yang et al., 2019). This reaction consumes protons, thereby increasing intracellular pH. The resulting shift in carbonate equilibrium favors the conversion of bicarbonate (HCO_3_^-^) to carbonate ions (CO_3_^2-^), which in turn promotes calcium carbonate precipitation. Conversely, when elemental sulfur is oxidized to sulfate, protons are released, decreasing intracellular pH and causing iACC dissolution. Thus, iACC function as a dynamic pH buffer: consuming protons through their dissolution and releasing them through their precipitation.

A similar hypothesis has been proposed for cyanobacteria, where photosynthesis elevates intracellular pH and thereby increases carbonate ion concentration, favoring iACC precipitation (Li et al., 2016). While these models hold for *Achromatium* and cyanobacteria, the present study shows that a proton-consuming primary metabolism is not strictly required for iACC biomineralization. In *Methylococcaceae*, aerobic methanotrophy does not consume protons, as shown by equation (1):

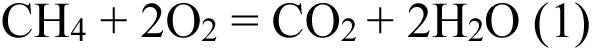

Moreover, the CO_2_ produced by aerobic methanotrophs should promote calcium carbonate dissolution as shown by Krause et al. (2014) and according to carbonate equilibria:

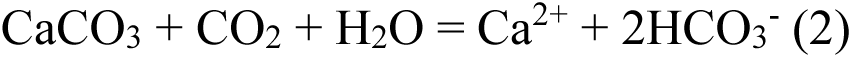

However, a portion of this CO_2_ may be diverted into small organic molecules via RuBiSCO-mediated carbon fixation, as shown in *Methylococcus*, and may therefore not react with CaCO_3_ (Oshkin et al., 2021). Overall, iACC precipitation in *Methylococcaceae* likely relies on an additional, secondary metabolic process—yet to be identified—that consumes protons and converts CO_2_ into carbonate ions so that:

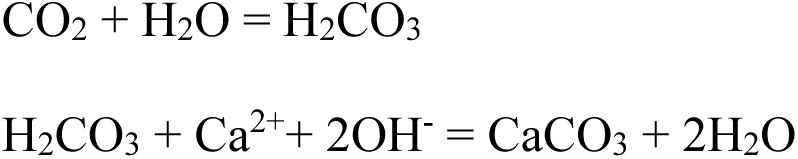

Alternatively, if CO_2_ fixation exceeds CO_2_ release from methanotrophy, the net CO_2_ production by the cell would become at least sometimes negative, shifting the carbonate equilibrium towards carbonate ions and favoring iACC precipitation. This hypothesis aligns with recent findings in magnetotactic iACC-forming *Alphaproteobacteria*, whose primary metabolism does not induce alkalinization but appears to support CO_2_ fixation (Mangin et al., 2025). Finally, calcium uptake is essential for storage within iACC. Ca^2+^/H^+^ protein exchangers (Cax proteins) are plausible candidates and we have detected *cax* genes in *Methylococcaceae* genomes (data not shown). However, these transporters are generally inferred to function in calcium efflux rather than uptake in bacteria (Waditee et al., 2004), which contradicts the requirements for iACC formation. Future studies are needed to elucidate the specific metabolic and transport processes enabling iACC biomineralization in *Methylococcaceae*. Aerobic methane-oxidizing bacteria are widespread in many aquatic systems and have a significant environmental and ecological role since they are key in the regulation of emissions of CH_4_, a potent greenhouse gas, and provide CH_4_-derived organic carbon compounds to surrounding heterotrophic microorganisms (Khanongnuch et al., 2022). How the capability to biomineralize iACC in some of these organisms may play an overlooked regulatory role in these important biogeochemical processes remains to be understood.

### 4.3 Evolutionary origin of iACC biomineralization in *Methylococcaceae*

The *ccyA* gene has an ancient origin within cyanobacteria, explaining its broad phylogenetic distribution across the phylum, with multiple secondary losses in modern lineages (Benzerara et al., 2022). As mentioned above, *ccyA* genes encode calcyanin proteins composed of a long, conserved C-terminal (GlyZip)_3_ domain and a short N-terminal domain, which exhibits variability across species. This difference between the two domains suggests that there is a strong selective pressure acting on the C-terminal domain, keeping it largely invariant, whereas the N-terminal domain appears much less constrained and free to vary. These contrasting evolutionary pressures have driven the diversification of several calcyanin families in cyanobacteria, each characterized by distinct N-terminal domains, termed CoBaHMA, X, Y, and Z. The X, Y, and Z types are restricted to specific clades, i.e., *Gloeomargarita*, *Fischerella* (and closely related genera), and *Microcystis*, respectively. In contrast, the CoBaHMA-type calcyanin occurs in several distantly related lineages and has therefore been proposed as the most ancient form of calcyanin (Benzerara et al., 2022).

The calcyanin sequences identified in *Methylococcaceae* have specific characteristics not found in the four cyanobacterial types but share clear sequence and structural similarities with the *Microcystis* (Z-type) calcyanins, which can be observed especially over the GlyZip1 motif of the (GlyZip)3 domain but also in the N-terminal domain. This is unexpected given the vast evolutionary distance between *Methylococcaceae* and all the cyanobacterial clades. Phylogenetic analysis strongly supports the relationship between the *Methylococcaceae* and cyanobacterial sequences (Figure 4). The most parsimonious explanation for this observation is that *Methylococcaceae* acquired *ccyA* through horizontal gene transfer (HGT) from a cyanobacterial ancestor closely related to *Microcystis*. Alternative explanations would be that (i) *ccyA* was present in the last common ancestor of *Methylococcaceae* and *Cyanobacteria*—two highly distant lineages—and subsequently lost independently thousands of times in all other bacterial lineages, or (ii) that *Methylococcaceae* and *Cyanobacteria* independently evolved (GlyZip)3-encoding genes with similar amino acid residues at key positions and independently linked them to small, variable N-terminal domains. However, both scenarios are far less parsimonious than the hypothesis of a single HGT event between *Methylococcaceae* and *Cyanobacteria*. The fact that the phylogeny of the *Methylococcaceae* calcyanin sequences mirrors that of the corresponding species (Figure 4), strongly suggests a single HGT acquisition by an ancient *Methylococcaceae* species, followed by subsequent divergence in several *Methylococcaceae* subgroups, including several independent losses in the contemporary *Methylococcaceae* lineages lacking *ccyA*. Such independent losses have also been proposed to explain the current distribution of *ccyA* in the contemporary cyanobacterial lineages (Benzerara et al., 2022).

About 1/3 of known *Microcystis* genomes contain *ccyA*, which was also likely ancestral in this genus and lost secondarily in several lineages (Gaetan et al., 2023). Consistently with the presence of *ccyA*, abundant iACC-forming *Microcystis* strains have been observed in several lakes (Gaetan et al., 2023). One plausible ecological setting for the HGT of *ccyA* from *Microcystis* to *Methylococacceae* is exemplified by Lake Cadagno, a meromictic lake (i.e., permanently stratified with an anoxic monimolimnion). At its chemocline, methane removal is driven by aerobic methane oxidizers which are physically associated with cyanobacteria (Milucka et al., 2015). More precisely, physical associations between *Microcystis* and diverse *Methylococcales* are common in lakes and ponds, where they may be responsible for an overlooked sink for methane (Li et al., 2021). Similarly, microbial consortia comprising cyanobacteria and methanotrophic bacteria are employed in oxygen-deficient wastewater treatment plants to remove methane (Safitri et al., 2021). In these consortia, cyanobacteria produce O_2_ that fuels aerobic methanotrophy, while the resulting CO_2_ is taken up by the phototrophs. Such tight metabolic coupling provides an ideal setting for close and sustained interactions between cyanobacteria and *Methylococcales*, potentially facilitating HGT. One of these HGT events appears to have transmitted the capability to biomineralize iACC from cyanobacteria to *Methylococcaceae*.

## Conclusions

This study shows that aerobic methanotrophs belonging to the *Methylococcaceae* family form iACC. They most likely acquired the *ccyA* gene, a diagnostic marker of iACC biomineralization in cyanobacteria, through HGT from an ancestral *Microcystis*-like cyanobacterial donor. This HGT event may have taken place in a permanently stratified lake or in any other anoxic-oxic transition zone in aquatic habitats, where *Microcystis* and *Methylococcaceae* can co-exist. *Methylococcaceae* forming iACC may be abundant in some environments. Yet, how iACC are formed and, in particular, how much of the carbon from methane oxidation is stored within iACC and diverted from the production of small organic carbon molecules by these bacteria remains to be quantified.

## Data Availability Statement

All gene sequences were retrieved from GTDB as indicated in the Material and Method section and no new sequence was generated by this study. Microscopy and spectroscopy data are available on Zenodo: https://doi.org/10.5281/zenodo.17846939

## Acknowledgements

This study was supported by the Agence Nationale de la Recherche (ANR), under grants Cyanoman, ANR-24-CE44-5543 and Carbomagnet ANR-21-CE01-0010 and the ERC grants Calcyan (PI: K. Benzerara, Grant Agreement no. 307110) and Plast-Evol (PI: D. Moreira, Grant Agreement no. 787904. We thank synchrotron ALBA (Barcelona, Spain) for providing beamtime on the cryo transmission X-ray microscopy at the Mistral beamline under proposal number 2023027368. We thank synchrotron SOLEIL (Saclay, France) for providing beamtime to conduct XANES measurements on the LUCIA beamline under proposal number 20220623. We also thank the IMPMC analytical platforms that enabled the data acquisition: Imène Estève for the SEM-EDXS support; Maxime Guillaumet and Keevin Beneut for IR spectroscopy support at IMPMC.

