## Supplementary figures and images for "Intracellular amorphous calcium carbonate biomineralization in methanotrophic gammaproteobacteria was acquired by horizontal gene transfer from cyanobacteria"

### Figure S1

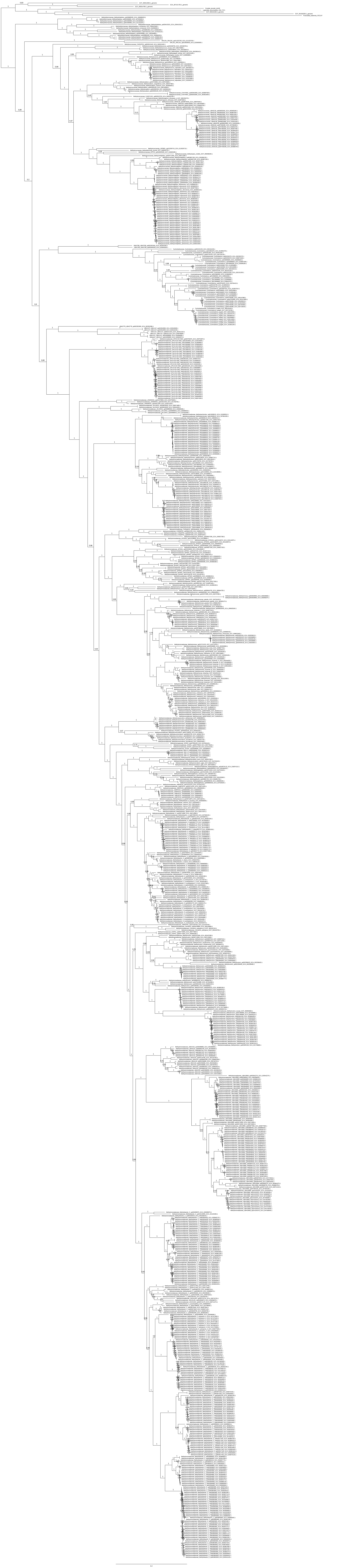
