## Supplementary material for "Intracellular amorphous calcium carbonate biomineralization in methanotrophic gammaproteobacteria was acquired by horizontal gene transfer from cyanobacteria": Supp Figures

**Supplementary Figures**

**Figure S1: see complete, un-collapsed tree relative to the main manuscript figure 1**

**Figure S2:** Comparison of the HCA plots of the *Microcystis* (represented by *M. aeruginosa* PCC 9443) and *Methylococcaceae* (represented by *M. geothermalis*) sequences, highlighting their overall similarity, including in their N-terminal regions, rich in alanine and poor in strong hydrophobic amino acids. The AF3 3D structure models of the two proteins illustrate that their N-terminal sequences (pink) are predicted to fold as long alpha-helices. However, despite these similarities, the sequences of the N-terminal regions cannot be aligned robustly. The protein amino acid sequences (one-letter code) are displayed on a duplicated alpha-helical net, on which the strong hydrophobic amino acids (V, I, L, F, M, Y, and W) are contoured. The latter form clusters, which mainly correspond to the internal faces of regular secondary structures (α-helices and β-strands). The way to read the primary (1D) and secondary (2D) structures is shown with arrows (one amino acid or one hydrophobic cluster after another, respectively), whereas special symbols used for four amino acids with specific structural properties (P, G, S, and T) are described in the inset.

**Time (hours)**

**OD_600nm_**

τ = 5.79 ±0.08 h

τ = 13.08 ±0.82 h

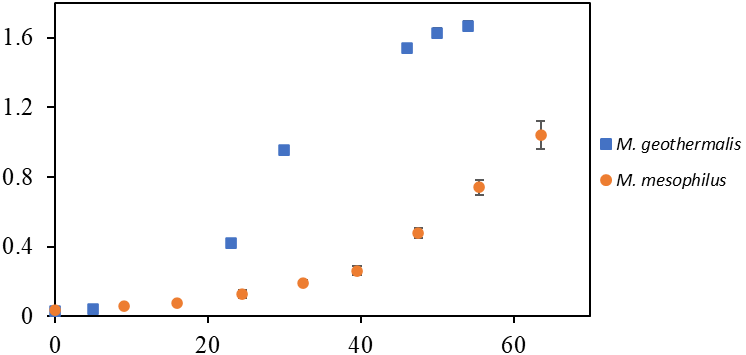

|  | M. geothermalis |
| --- | --- |
| 0 | 0.0303333 |
| 5 | 0.0393333 |
| 23 | 0.421333 |
| 30 | 0.953333 |
| 46 | 1.54 |
| 50 | 1.63 |
| 54 | 1.67 |
| 118 | 1.61333 |

|  | M. mesophilus |
| --- | --- |
| 0 | 0.035 |
| 9 | 0.055 |
| 16 | 0.0765 |
| 24.5 | 0.1275 |
| 32.5 | 0.1925 |
| 39.5 | 0.2615 |
| 47.5 | 0.47875 |
| 55.5 | 0.74 |
| 63.5 | 1.043 |

**Figure S3**: Growth curves of *M. geothermalis* (squares) and *M. mesophilus* (circles). Cultures were conducted in triplicates. Optical density at 600 nm was used as a proxy for cell abundances. Error bars indicates standard deviation. When not visible, error bars are smaller than the symbol size. Generation times (t) are provided for both strains.

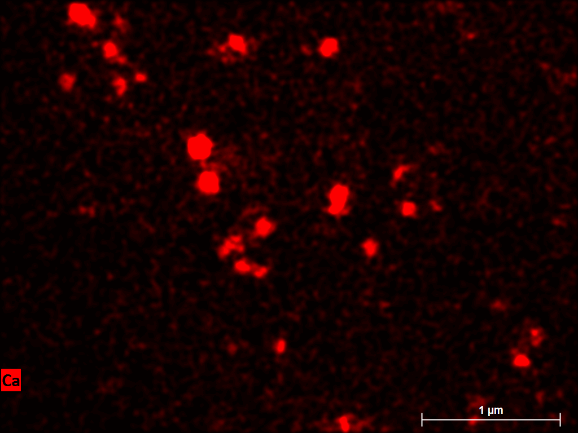

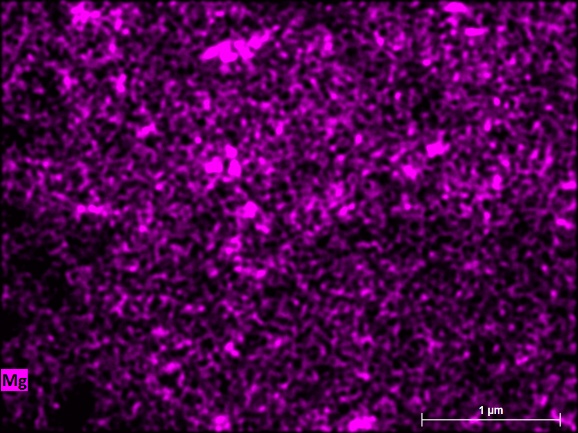

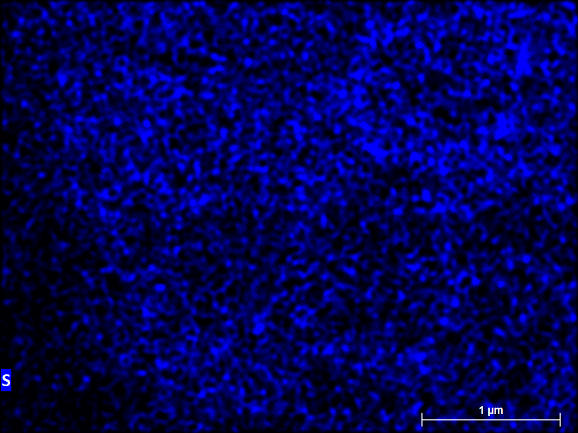

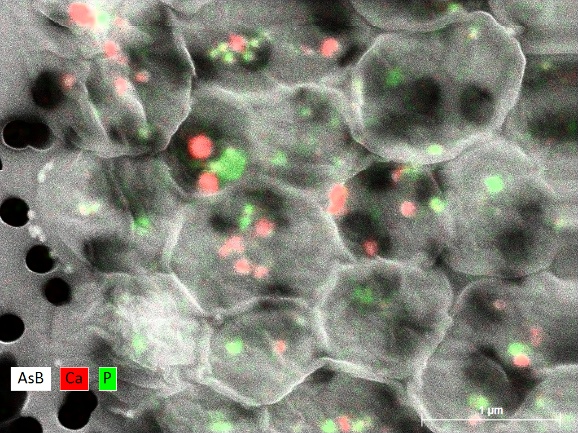

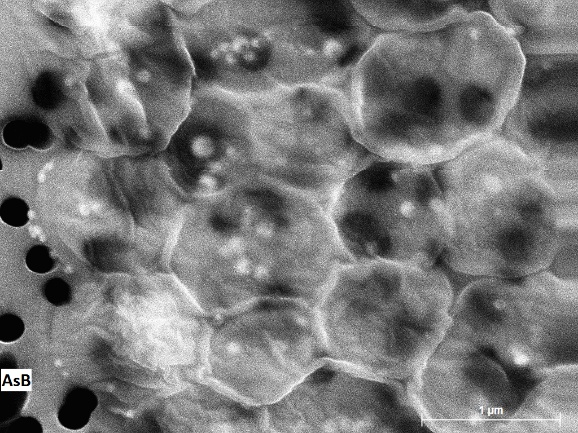

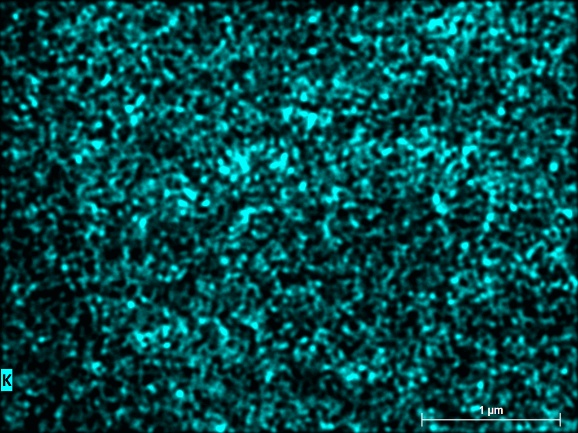

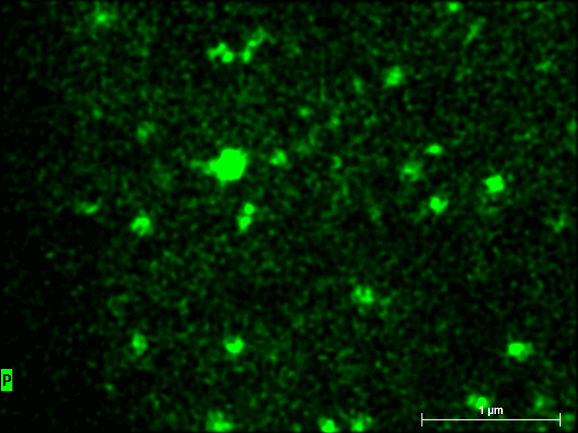

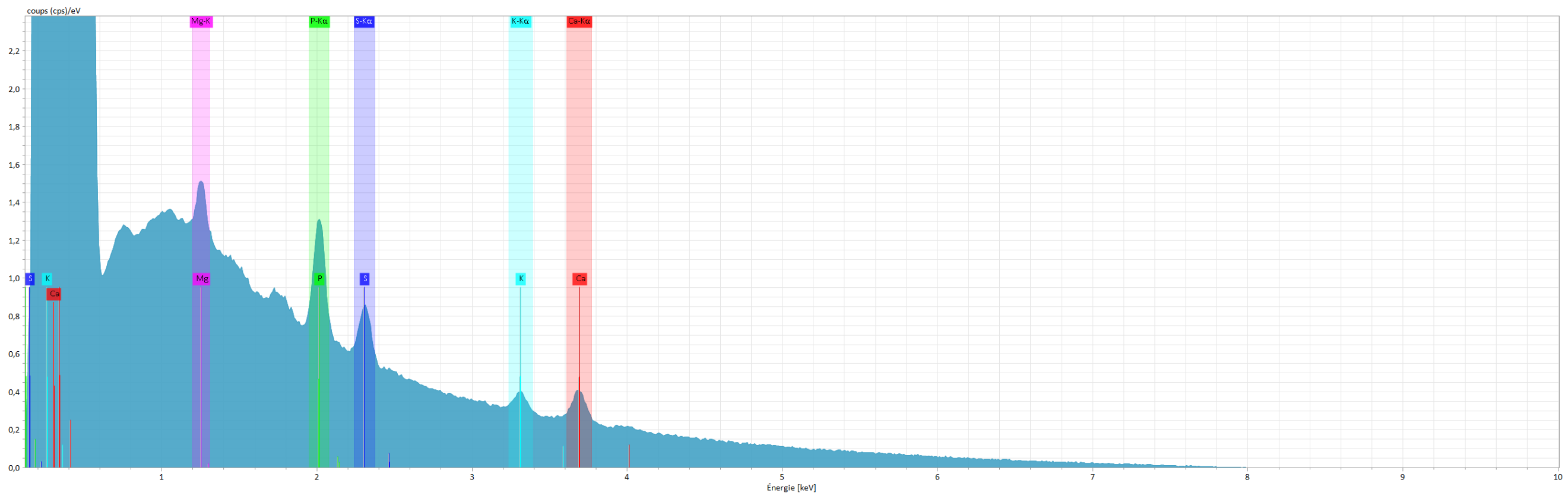

**Figure S4**: Scanning electron microscopy analyses of *Methylococcus geothermalis*. SEM image, overlay of SEM image with Ca and P maps, Ca, P, K, Mg and S maps as well as the spectrum of the whole scanned area are provided. Raw data are available on Zenodo: <https://doi.org/10.5281/zenodo.17846939>

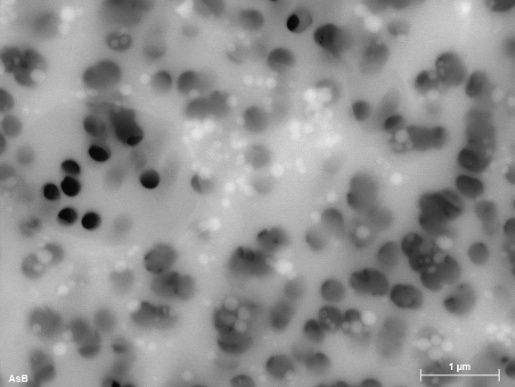

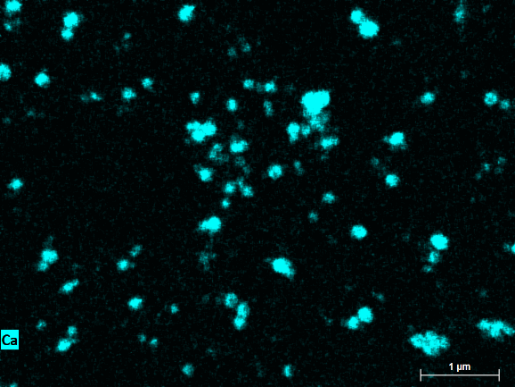

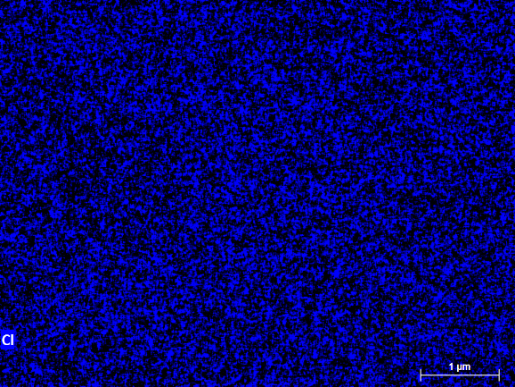

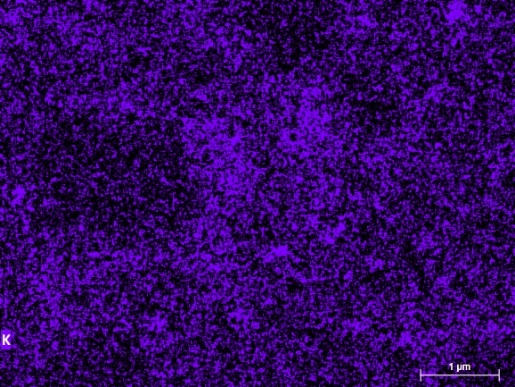

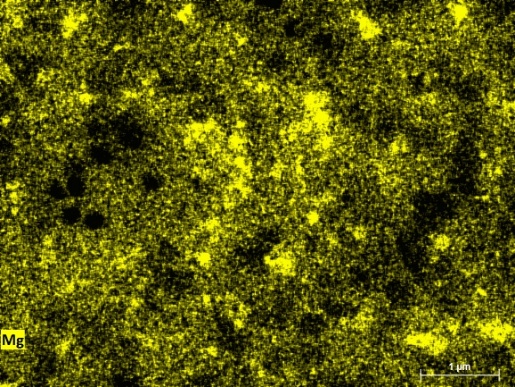

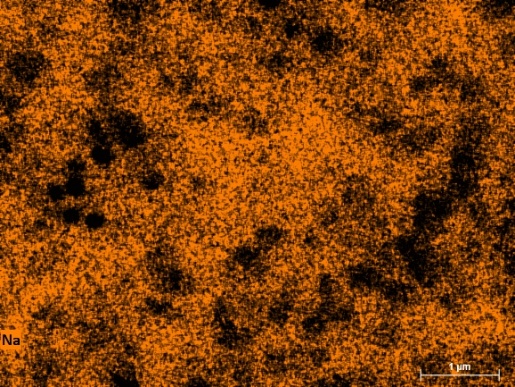

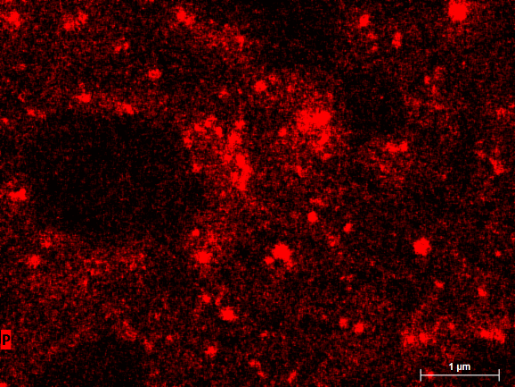

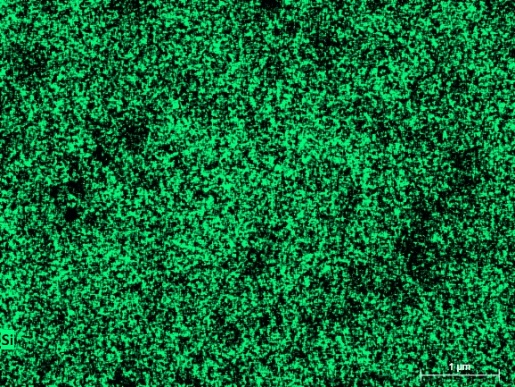

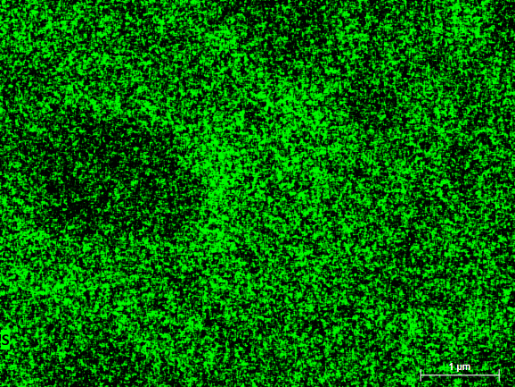

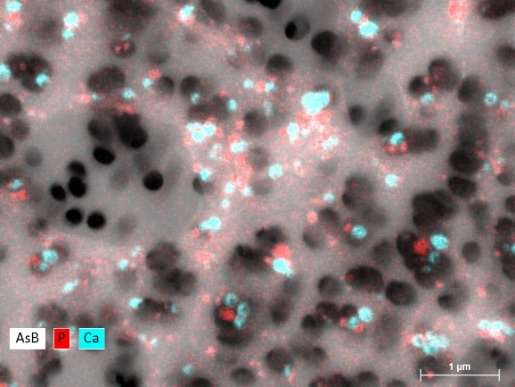

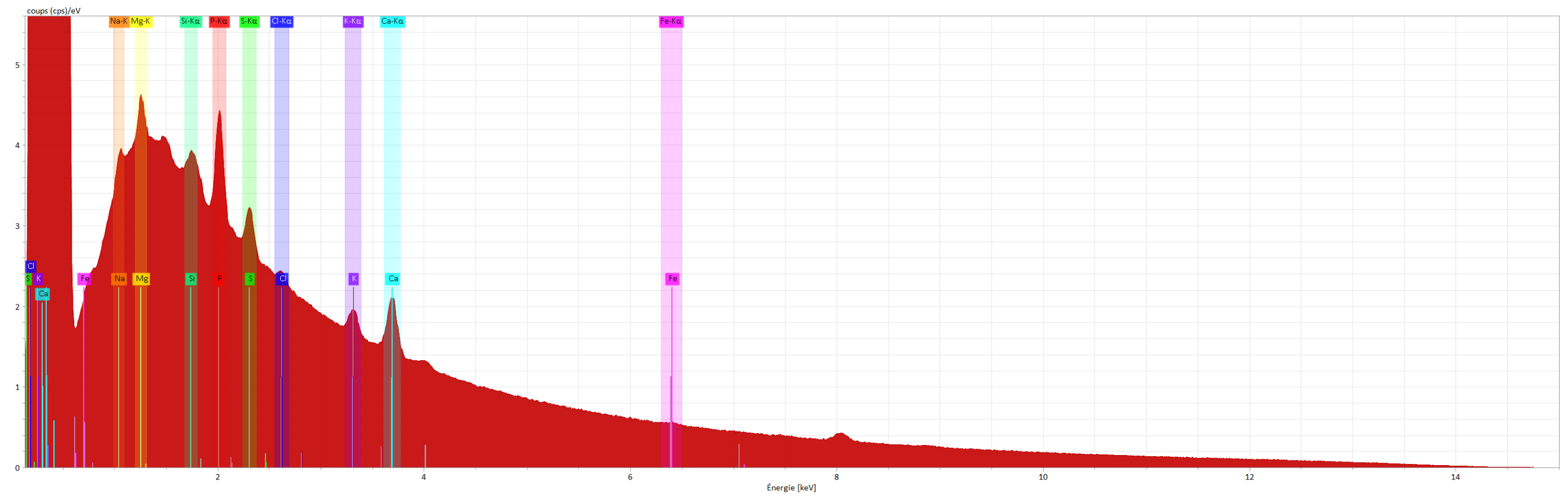

**Figure S5**: Scanning electron microscopy analyses of *Methylococcus mesophilus*. SEM image, overlay of SEM image with Ca and P maps, Ca, P, K, S, Mg, Cl, Si and Na maps as well as the spectrum of the whole scanned area are provided. Raw data are available on Zenodo: <https://doi.org/10.5281/zenodo.17846939>

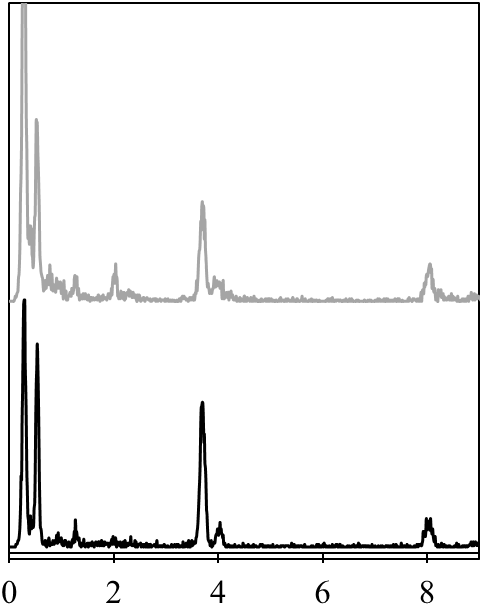

Energy (keV)

X-ray intensity

Cu Kα

Ca Kβ

Mg Kα

Ca Kα

C Kα

O Kα

P Kα

*M. mesophilus*

*M. geothermalis*

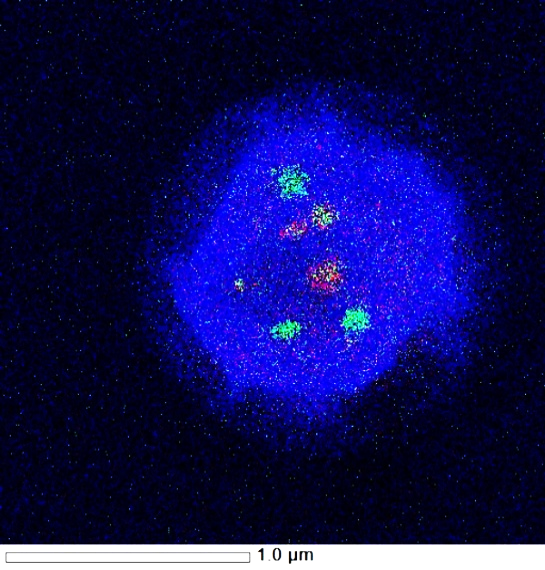

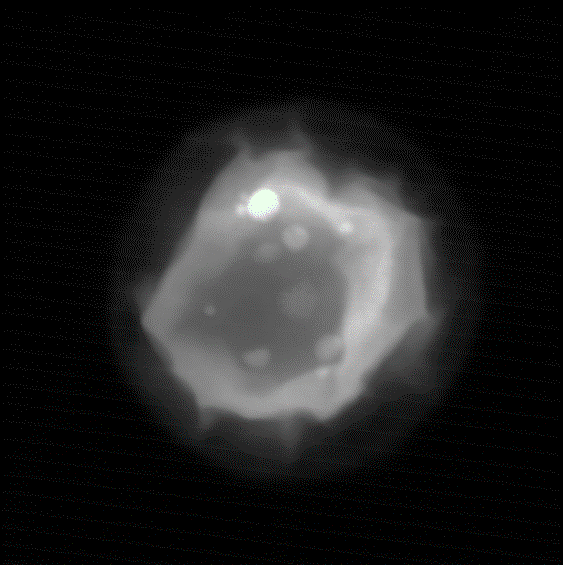

500 nm

500 nm

*M. mesophilus*

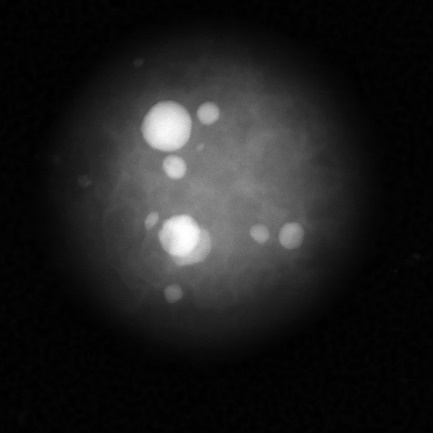

500 nm

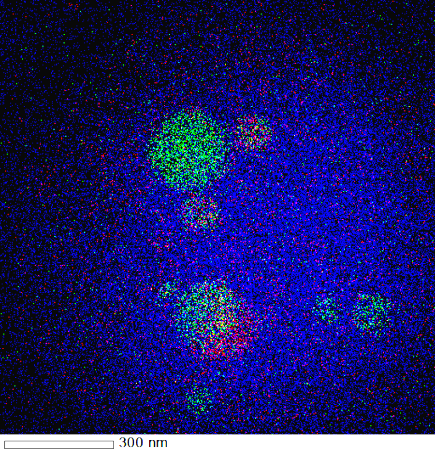

300 nm

*M. geothermalis*

A

B

C

D

E

|  | geothermalis | mesophilus 2 |
| --- | --- | --- |
| 0 | 0 | 190 |
| 0.01 | 0 | 190 |
| 0.02 | 0 | 190 |
| 0.03 | 0 | 190 |
| 0.04 | 0 | 190 |
| 0.05 | 0 | 190 |
| 0.06 | 0 | 190 |
| 0.07 | 0 | 190 |
| 0.08 | 0 | 190 |
| 0.09 | 0 | 191 |
| 0.1 | 0 | 192 |
| 0.11 | 0 | 194 |
| 0.12 | 2 | 193 |
| 0.13 | 2 | 194 |
| 0.14 | 2 | 196 |
| 0.15 | 3 | 197 |
| 0.16 | 5 | 198 |
| 0.17 | 5 | 208 |
| 0.18 | 7 | 211 |
| 0.19 | 11 | 220 |
| 0.2 | 19 | 242 |
| 0.21 | 32 | 274 |
| 0.22 | 55 | 288 |
| 0.23 | 52 | 323 |
| 0.24 | 98 | 397 |
| 0.25 | 122 | 443 |
| 0.26 | 162 | 481 |
| 0.27 | 188 | 502 |
| 0.28 | 191 | 503 |
| 0.29 | 191 | 496 |
| 0.3 | 140 | 468 |
| 0.31 | 123 | 392 |
| 0.32 | 94 | 331 |
| 0.33 | 59 | 308 |
| 0.34 | 42 | 269 |
| 0.35 | 24 | 239 |
| 0.36 | 21 | 249 |
| 0.37 | 12 | 239 |
| 0.38 | 10 | 234 |
| 0.39 | 18 | 237 |
| 0.4 | 24 | 248 |
| 0.41 | 16 | 244 |
| 0.42 | 14 | 246 |
| 0.43 | 16 | 226 |
| 0.44 | 19 | 224 |
| 0.45 | 15 | 216 |
| 0.46 | 17 | 222 |
| 0.47 | 30 | 240 |
| 0.48 | 34 | 255 |
| 0.49 | 56 | 257 |
| 0.5 | 96 | 294 |
| 0.51 | 125 | 330 |
| 0.52 | 138 | 326 |
| 0.53 | 157 | 328 |
| 0.54 | 141 | 305 |
| 0.55 | 121 | 294 |
| 0.56 | 93 | 277 |
| 0.57 | 54 | 256 |
| 0.58 | 31 | 226 |
| 0.59 | 15 | 220 |
| 0.6 | 14 | 213 |
| 0.61 | 11 | 211 |
| 0.62 | 5 | 209 |
| 0.63 | 3 | 208 |
| 0.64 | 2 | 203 |
| 0.65 | 3 | 198 |
| 0.66 | 2 | 202 |
| 0.67 | 3 | 199 |
| 0.68 | 5 | 206 |
| 0.69 | 1 | 205 |
| 0.7 | 1 | 200 |
| 0.71 | 0 | 205 |
| 0.72 | 2 | 207 |
| 0.73 | 5 | 210 |
| 0.74 | 2 | 203 |
| 0.75 | 7 | 205 |
| 0.76 | 5 | 203 |
| 0.77 | 1 | 218 |
| 0.78 | 2 | 211 |
| 0.79 | 3 | 207 |
| 0.8 | 3 | 199 |
| 0.81 | 2 | 211 |
| 0.82 | 2 | 202 |
| 0.83 | 5 | 198 |
| 0.84 | 3 | 203 |
| 0.85 | 2 | 201 |
| 0.86 | 4 | 198 |
| 0.87 | 5 | 201 |
| 0.88 | 3 | 197 |
| 0.89 | 4 | 201 |
| 0.9 | 9 | 208 |
| 0.91 | 4 | 200 |
| 0.92 | 3 | 205 |
| 0.93 | 11 | 205 |
| 0.94 | 4 | 201 |
| 0.95 | 7 | 200 |
| 0.96 | 5 | 203 |
| 0.97 | 4 | 205 |
| 0.98 | 6 | 204 |
| 0.99 | 7 | 196 |
| 1 | 2 | 200 |
| 1.01 | 3 | 197 |
| 1.02 | 1 | 197 |
| 1.03 | 3 | 192 |
| 1.04 | 1 | 206 |
| 1.05 | 5 | 196 |
| 1.06 | 5 | 194 |
| 1.07 | 5 | 194 |
| 1.08 | 3 | 198 |
| 1.09 | 3 | 191 |
| 1.1 | 1 | 192 |
| 1.11 | 0 | 191 |
| 1.12 | 2 | 194 |
| 1.13 | 4 | 194 |
| 1.14 | 3 | 194 |
| 1.15 | 3 | 196 |
| 1.16 | 0 | 197 |
| 1.17 | 3 | 194 |
| 1.18 | 0 | 194 |
| 1.19 | 4 | 200 |
| 1.2 | 5 | 200 |
| 1.21 | 6 | 196 |
| 1.22 | 3 | 198 |
| 1.23 | 8 | 199 |
| 1.24 | 12 | 208 |
| 1.25 | 4 | 210 |
| 1.26 | 21 | 209 |
| 1.27 | 11 | 203 |
| 1.28 | 12 | 201 |
| 1.29 | 10 | 209 |
| 1.3 | 6 | 195 |
| 1.31 | 7 | 196 |
| 1.32 | 5 | 200 |
| 1.33 | 7 | 194 |
| 1.34 | 2 | 191 |
| 1.35 | 1 | 195 |
| 1.36 | 3 | 194 |
| 1.37 | 0 | 192 |
| 1.38 | 3 | 195 |
| 1.39 | 2 | 193 |
| 1.4 | 1 | 192 |
| 1.41 | 1 | 196 |
| 1.42 | 6 | 194 |
| 1.43 | 2 | 192 |
| 1.44 | 2 | 192 |
| 1.45 | 0 | 196 |
| 1.46 | 2 | 193 |
| 1.47 | 2 | 193 |
| 1.48 | 2 | 192 |
| 1.49 | 3 | 191 |
| 1.5 | 1 | 192 |
| 1.51 | 0 | 195 |
| 1.52 | 3 | 191 |
| 1.53 | 2 | 192 |
| 1.54 | 3 | 190 |
| 1.55 | 2 | 193 |
| 1.56 | 1 | 193 |
| 1.57 | 1 | 193 |
| 1.58 | 4 | 191 |
| 1.59 | 1 | 191 |
| 1.6 | 2 | 193 |
| 1.61 | 2 | 192 |
| 1.62 | 0 | 192 |
| 1.63 | 3 | 191 |
| 1.64 | 2 | 192 |
| 1.65 | 4 | 192 |
| 1.66 | 3 | 193 |
| 1.67 | 2 | 193 |
| 1.68 | 1 | 191 |
| 1.69 | 2 | 194 |
| 1.7 | 4 | 195 |
| 1.71 | 4 | 193 |
| 1.72 | 1 | 194 |
| 1.73 | 2 | 190 |
| 1.74 | 4 | 192 |
| 1.75 | 1 | 191 |
| 1.76 | 5 | 193 |
| 1.77 | 4 | 194 |
| 1.78 | 1 | 191 |
| 1.79 | 2 | 195 |
| 1.8 | 1 | 193 |
| 1.81 | 1 | 193 |
| 1.82 | 4 | 192 |
| 1.83 | 2 | 192 |
| 1.84 | 2 | 193 |
| 1.85 | 2 | 192 |
| 1.86 | 3 | 191 |
| 1.87 | 2 | 193 |
| 1.88 | 3 | 194 |
| 1.89 | 2 | 196 |
| 1.9 | 0 | 195 |
| 1.91 | 3 | 196 |
| 1.92 | 1 | 192 |
| 1.93 | 3 | 194 |
| 1.94 | 3 | 194 |
| 1.95 | 5 | 200 |
| 1.96 | 6 | 195 |
| 1.97 | 2 | 204 |
| 1.98 | 8 | 209 |
| 1.99 | 6 | 207 |

**Figure S6**: Scanning transmission electron microscopy (STEM) analyses of cultivated *Methylococcaceae* in the high angle annular darkfield mode (HAADF). (A-B) STEM-HAADF image of a *Methylococcus mesophilus* cell and corresponding overlay of EDXS maps of carbone (blue), calcium (green) and phosphorus (red). The green objects are rich in Ca, poor in P and interpreted as iACC inclusions, while the red objects are polyphosphate inclusions. (C-D) Same for *Methylococcus geothermalis*. (E) EDXS spectra of iACC in *M. mesophilus* (top) and *M. geothermalis* (bottom). X-ray emission lines are indexed. Some P is detected in the *M. mesophilus* iACC spectrum but is likely due to overlap of iACC granules and PolyP.

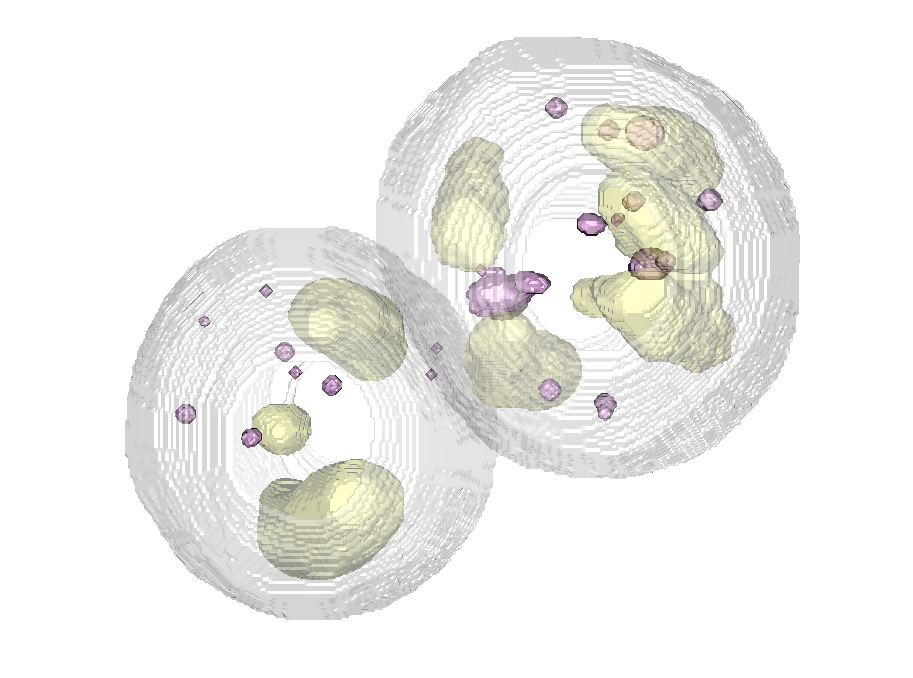

y

x

**B**

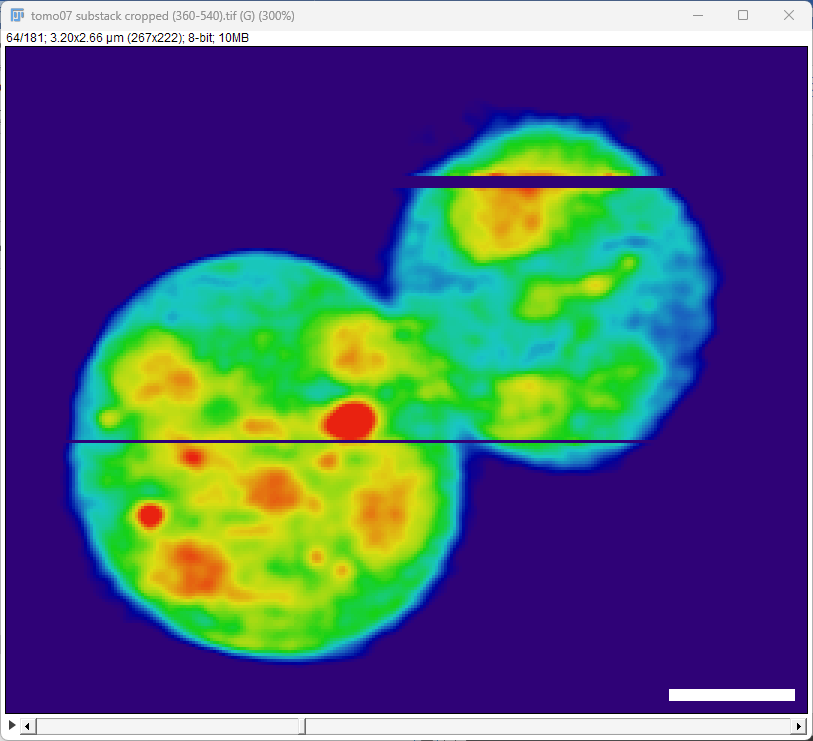

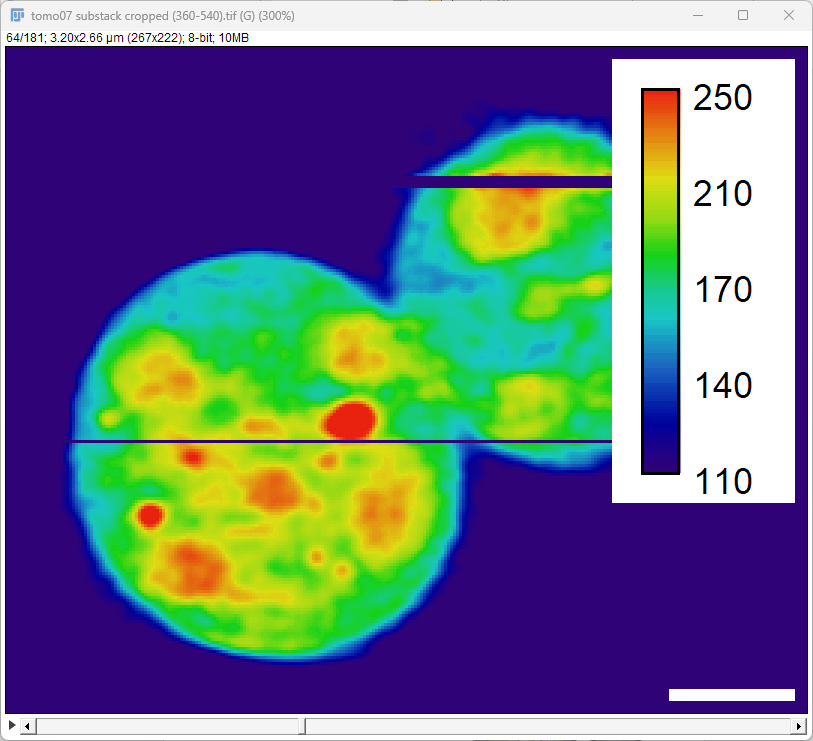

y

x

**A**

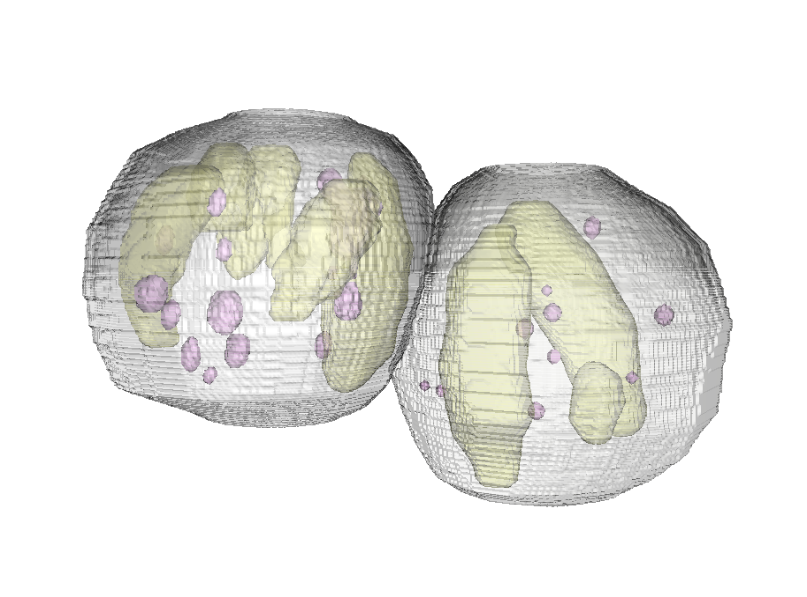

z

y

**Figure S7**: Cryo-soft X-ray tomography (c-SXT) of two ice-vitrified *Methylococcus geothermalis* cells (distinct from the two ones shown in Figure 7). The image intensity corresponds to X-ray absorption by intracellular material at 520 eV (i.e. below the O K-edge). (A) Virtual mid-cell z-slice of two *M. geothermalis* cells in reconstructed tomogram. Scale bar is 500 nm. (B) Volume reconstruction of cells in (A) shown in two perspectives with Ca-rich granules in pink, thylakoid/phosphate inclusions in beige and outer cell membrane in grey.
