## Supplementary material for "Intracellular amorphous calcium carbonate biomineralization in methanotrophic gammaproteobacteria was acquired by horizontal gene transfer from cyanobacteria": Table S3

|  | ***Methylococcus geothermalis*** | ***Microcystis aeruginosa***  ***PCC_9443*** | ***Fischerella***  ***sp_NIES-4106*** | ***Synechococcus_***  ***sp_PCC_6312*** | ***Gloeomargarita lithophora***  ***Alchichica*** |
| --- | --- | --- | --- | --- | --- |
| ***Methylococcus geothermalis*** | 1/2: 1.46 Å (38 aa)  1/3: 1.83 Å (43 aa)  2/3: 1.60 Å (42 aa) | **1/1: 1.49 Å (66 aa) TM 0.84,61% id**  2/2: 1.39 Å (39 aa)  3/3: 1.56 Å (52 aa) | **1/1: 2.06 Å (66 aa)**  **TM 0.87,31% id**  2/2: 1.90 Å (45 aa)  3/3: 1.75 Å (54 aa) | **1/1: 1.92 Å (66 aa)**  **TM 0.82,26% id**  2/2: 1.31 Å (39 aa)  3/3: 2.33 Å (53 aa) | **1/1:1.67 Å (59 aa)**  **TM0.76,39% id**  2/2: 1.46 Å (39 aa)  3/3: 1.82 Å (54 aa) |
| ***Microcystis aeruginosa***  ***PCC_9443*** |  | 1/2: 1.81 Å (58 aa)  1/3: 1.46 Å (42 aa)  2/3: 1.79 Å (47 aa) | **1/1: 1.90 Å (65 aa)**  **TM 0.88, 34% id**  2/2: 1.68 Å (60 aa)  3/3: 1.70 Å (53 aa) | **1/1: 2.06 Å (63 aa)**  **TM 0.81, 35% id**  2/2: 1.68 Å (51 aa)  3/3: 2.08 Å (40 aa) | **1/1: 1.55 Å (57 aa)**  **TM 0.77, 45% id**  2/2: 1.55 Å (51 aa)  3/3: 0.76 Å (53 aa) |
| ***Fischerella***  ***sp_NIES-4106*** |  |  | 1/2: 1.28 Å (63 aa)  1/3: 1.60 Å (46 aa)  2/3: 2.35 Å (44 aa) | **1/1: 1.73 Å (70 aa)**  **TM 0.79, 36% id**  2/2: 1.98 Å (50 aa)  3/3: 2.06 Å (45 aa) | **1/1: 1.29 Å (63 aa)**  **TM 0.75, 39% id**  2/2: 1.67 Å (52 aa)  3/3: 1.76 Å (54 aa) |
| ***Synechococcus_***  ***sp_PCC_6312*** |  |  |  | 1/2: 1.46 Å (51 aa)  1/3: 1.22 Å (43 aa)  2/3: 1.31 Å (41 aa) | **1/1: 2.04 Å (58 aa)**  **TM 0.70, 40% id**  2/2: 1.07 Å (56 aa)  3/3: 2.09 Å (41 aa) |
| ***Gloeomargarita lithophora Alchichica*** |  |  |  |  | 1/2: 1.30 Å (54 aa)  1/3: 1.52 Å (48 aa)  2/3: 1.66 Å (53 aa) |

**Table S3:** **Metrics related to the superimpositions of the GlyZip 3D structure models.** Root Mean Square Deviation (RMSD) values are indicated, as well as the number of aligned amino acids (aa, C-alpha atoms). For the comparisons of GlyZip motif 3D structure models within a species (diagonal) and between species (GlyZip2 and GlyZip3), only the GlyZip conserved core can be superimposed, while the N- and C-terminal extremities are variable. Only the GlyZip1 3D structure models can be superimposed over their whole length (highlighted in yellow and grey, with TM scores and sequence identities). The *Methyloccocus geothermalis* and *Microcystis aeruginosa* GlyZip1 superimposition (yellow) shows the lowest RMSD and highest sequence identity.
