## Supplementary material for "Intracellular amorphous calcium carbonate biomineralization in methanotrophic gammaproteobacteria was acquired by horizontal gene transfer from cyanobacteria": Table S4

Table Sx: Wavenumbers and band assignment for characteristics vibrations found in the mid IR spectra of bacteria. The band assignment was achieved based on Mehta et al. (2022).

| **Band Index** | **Wavenumbers (cm^-1^)** | **Main functional group** |
| --- | --- | --- |
| 1 | 3279 | water |
| 2 | 2856-2960 | Methyl and methylene group vibrations from lipids and fatty acids |
| 3 | 1641 | Amide I from proteins |
| 4 | 1528 | Amide II from proteins |
| 5 | 1444 | Methyl groups of lipids and C-O group in carboxylic acids |
| 6 | 1391 | Methyl groups of lipids and C-O group in carboxylic acids |
| 7 | 1241 | Nucleic acids, phosphoryl groups |
| 8 | 1153 | P=O group of nucleic acids and/or polyphosphates and C-O vibrations from polysaccharides |
| 9 | 1029-1058 | P=O group of nucleic acids and/or polyphosphates and C-O vibrations from polysaccharides |
| 10 | 919 | P=O group of nucleic acids and/or polyphosphates and C-O vibrations from polysaccharides |
| 11 | 860 | Out-of-plane bending of carbonates in ACC (ν_2_) |
